# Notch mediated lateral inhibition is shaped by morphological differences to reinforce bias toward signal-sending or receiving roles

**DOI:** 10.64898/2026.03.27.712410

**Authors:** Prachi Richa, Charalambos Roussos, Chengxi Zhu, Martin O. Lenz, Shahar Kasirer, David Sprinzak, Sarah Bray

## Abstract

During neurogenesis, neuroblasts are selected from proneural-competent cells through lateral inhibition, a process controlled by the evolutionarily conserved Notch signalling pathway. By tracking transcription from Notch-target genes and cell morphologies in real time, we discovered that the presumptive neuroblast never initiates target-gene transcription. This implies a pre-existing bias directs Notch signalling. The bias correlates with a heterogeneity in apical cell areas which is further reinforced during neuroblast selection. Additionally, the length and duration of neuroblast-neighbour cell contacts predict the likelihood of transcription. Using mathematical modelling we show that lateral inhibition seeded with subtle morphological differences can bias cells toward signal-sending or receiving roles before transcriptional feedback occurs. Notch activation further alters apical cell area, reinforcing the initial bias. We propose that signalling and cell mechanics work together to ensure the robust selection of a single neural precursor.

## Introduction

The selection of neural precursor cells from a pool of cells with proneural competence is regulated by activity of the evolutionarily conserved Notch signalling pathway, a process called lateral inhibition^1^. One feature of lateral inhibition models is an intercellular transcriptional feedback regulatory loop^2–4^. This involves cell-fate promoting factors, such as proneural proteins of the Achaete-Scute family, upregulating the expression of Notch ligands. Notch activation results in the expression of HES gene family repressors, *Enhancer of split* [*E(spl)*] genes in *Drosophila*, which suppress activity of proneural proteins and, hence, that of the ligands. This feedback mechanism can amplify small initial differences, resulting in an all-or-none switch where one cell retains proneural activity and becomes the neural stem cell, whereas its neighbours are prevented from doing so. While there is evidence to support many aspects of this feedback mechanism, several observations challenge our understanding of how this operates in the real time of many developmental fate decisions^5,6^. Notably it is unclear whether all cells have equal potential to become neural and rely on a stochastic event to tip the balance in favour of one cell retaining neural determinants, as originally envisaged, or whether there is a pre-existing bias on which lateral inhibition operates.

It is also evident that, while transcriptional feedback is important, regulation of Delta and Notch can occur on multiple levels. For example, selection of neural precursors is largely normal under conditions where uniform Notch or Delta expression is provided, arguing that other modes of regulation must be deployed^7–9^. Mechanisms that regulate Delta activity more directly, including the E3 ubiquitin ligase Neuralized, likely participate in the molecular switch^10–12^. In addition, cis-inhibition, inhibitory interactions between ligands and receptors present on the same cell surface^13–15^, modulates Notch activity levels in many contexts including adult sensory organs, where it sets a baseline of activity for the selection of a single precursor^15^. Finally, properties of contacts between cells can affect levels and durations of cell-to-cell signalling^16,17^.

Signalling is normally initiated when transmembrane ligands on adjacent cells interact with Notch receptors at the membrane^18^. The ligand exerts a mechanical pulling force, likely generated by it’s endocytosis, that brings about a series of proteolytic cleavages leading to release of the Notch intracellular domain, NICD^19^. Released NICD translocates into the nucleus and forms activation complexes that drive transcription of target genes including those of the HES gene family^20^. The mechanical nature of the activation process means that effective signalling may depend on physical properties like tissue stiffness, junctional tensions, intracellular contractility^21^. For example, delamination of cardiomyocytes during cardiac trabeculation elicits high levels of Notch activity in neighbours^22^. It’s proposed that this signalling is mechanics-induced and that Notch activity suppresses the actomyosin machinery to limit delamination. As Neuroblast (NB) selection during *Drosophila* embryogenesis unfolds within a dynamically remodelling neuroepithelium and the NB itself undergoes apical constriction and Myosin-II–dependent ingression^23,24^, it’s possible that similar mechanical regulation contributes to Notch-mediated lateral inhibition during this process. However, whether there is an interplay between morphogenetic changes and spatiotemporal signalling within proneural clusters remains unknown.

To investigate Notch signalling dynamics during lateral inhibition, we tracked real-time transcription from Notch-target genes^25,26^ in combination with quantitative morpho-dynamic measurements of cell properties. We coupled this with laser ablations and genetic manipulations to perturb different cells and the contacts between them. Strikingly, we find that there is a pre-existing bias such that the presumptive NB never initiates target-gene transcription. This correlates with a pre-existing heterogeneity in the apical areas that are reinforced during NB selection. Our observations suggest that lateral inhibition is initiated from a landscape in which subtle differences bias the cells towards signal-sending or signal-receiving roles before transcriptional feedback. Incorporating the dynamic area heterogeneities into a mathematical model of lateral inhibition is sufficient to replicate the patterns of transcription observed. We propose that lateral inhibition involves a coupled system in which signalling and cell mechanics act together within the proneural cluster to ensure robust selection of a single neural precursor.

## Results

### Notch-responsive transcription reveals an early bias within the proneural clusters

Each neuroblast (NB), originates from a proneural cluster, epithelial cells with neural potential conferred by expression of proneural proteins such as Achaete (Figure 1A, S1A)^27,28^. To investigate whether all cells in a cluster initially signal equivalently to one another, before one acquires a higher signalling potential, or whether an initial bias pre-patterns signalling, we set out to track Notch-dependent transcriptional dynamics in real time within proneural clusters. We introduced MS2 loops into endogenous Notch target genes by CRISPR engineering, producing *E(spl)m8–MS2* flies^26^. When combined with fluorescent MS2 binding protein, MCP::GFP, discrete transcription foci could be detected within nuclei. Their intensity is a direct measure of the number of RNA molecules being transcribed (Figure 1B)^29^. We focussed on the first S1 wave (stage 7–8) of neurogenesis which initiates just after mesoderm invagination is completed (Figure 1B; S1A)^27^. By including a membrane-marker (Spider-GFP) delaminating NBs could be identified by their reduction in apical area (Figure 1C).

**Figure 1.**
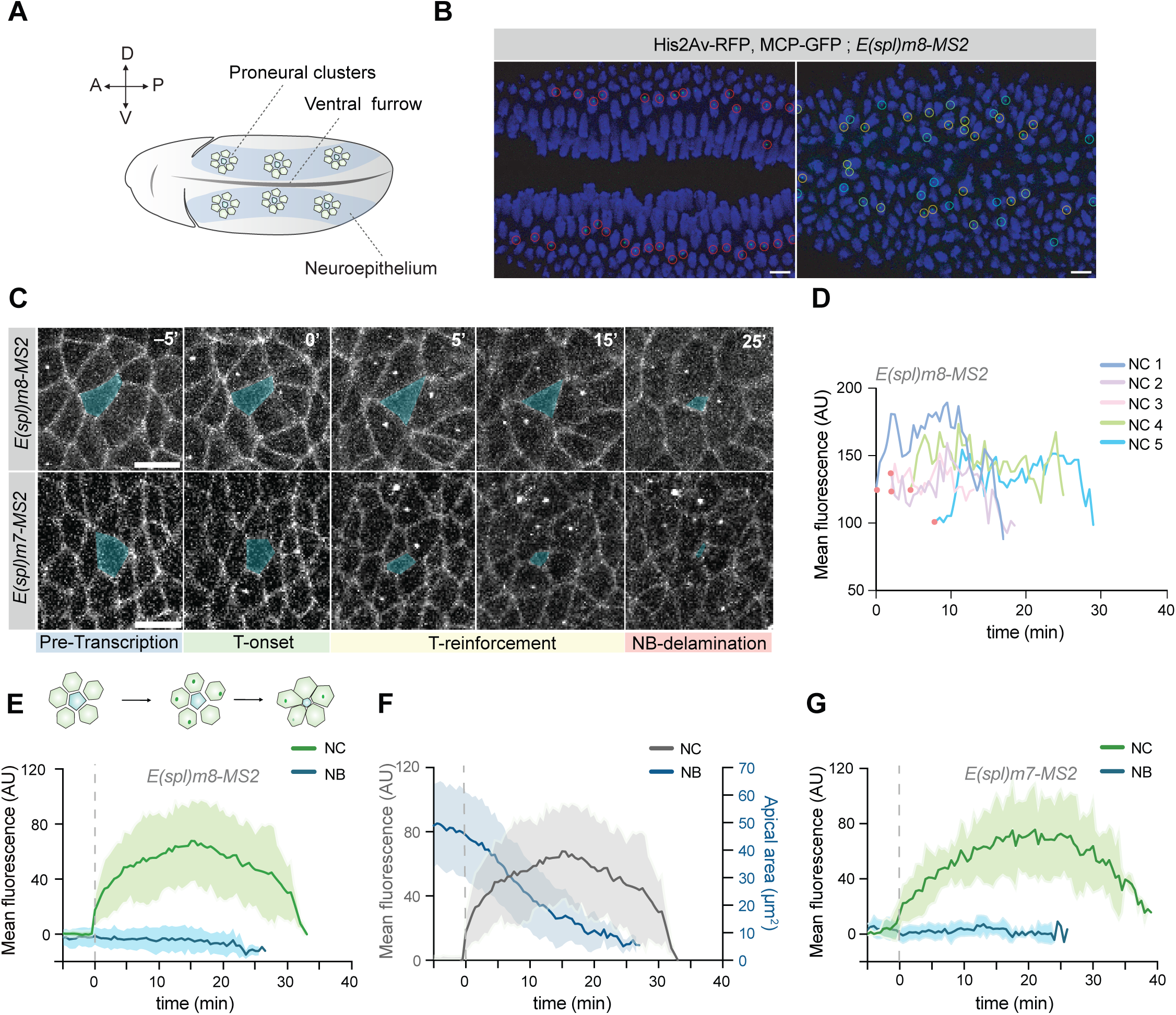
Tracking Notch-responsive transcription in proneural clusters reveals an early transcriptional bias. **(A)** Schematic illustrating proneural clusters within ventral neuroectoderm (blue shading) flanking the ventral furrow in stage 7–8 *Drosophila* embryo. Dorsal(D), ventral (V), Anterior (A) and posterior (P) axis are indicated. **(B)** Images from in vivo movie, *E(spl)m8–MS2* transcription is detected as puncta of MCP::GFP (green) within nuclei (blue, H2Av-RFP). Left image, mesectoderm (red circles) prior to ventral closure, right image, ventral neuroectoderm (yellow and blue circles). Scale bars, 10µm. **(C)** Time-lapse images from individual proneural clusters transcribing *E(spl)m8–MS2* or *E(spl)m7–MS2* in embryos expressing Spider-GFP to mark cell membranes. Presumptive neuroblast (NB) is shaded in cyan, identified by back-tracking from delamination (25’ right hand side). Transcription is aligned by time when *E(spl)m8/m7-MS2/*MCP puncta first detected (T-Onset 0’). Images at times relative to onset (0’) as indicated with NB-delamination at 25 min. Scale bars, 10µm. **(D)** *E(spl)m8-MS2*/MCP intensity profiles from individual cells within a representative cluster. Onset is asynchronous and amplitudes vary. **(E)** Mean fluorescence intensity of *E(spl)m8-MS2/MCP* aligned by transcription onset within a cluster (time 0). Puncta are detected in NCs (green), with an average duration per cluster of 25.90 ± 4.9 mins (mean ± SD). Presumptive NBs (blue) are transcriptionally silent. N=30 clusters, 84 NCs. **(F)** Comparison between *E(spl)m8–MS2* transcription per cluster (grey, replicated from E) and NB apical area (blue). Average duration of apical NB delamination, 18.43 ± 4.9 mins. **(G)** Mean fluorescence intensity of *E(spl)m7-MS2*/MCP aligned to transcription onset (time 0) as in E. *E(spl)m7–MS2* exhibits similar dynamics (average cluster duration, 32.23 ± 5.6 mins) and absence of activity in NBs. N=11 clusters, 34 NCs.

To characterize transcription dynamics at single-cell resolution, we performed 30–40-minute time-lapse imaging and extracted intensity of transcription foci in all tracked cells from identified clusters (Figure 1C, Video S1). First, *E(spl)m8–MS2* traces were aligned by onset of any detectable transcription within each cluster (time 0). We then compared the times at which foci first appeared and found that transcription initiated asynchronously within clusters (Figure 1C,D; S1B), with lag times of ∼5–7 minutes between nuclei. Once initiated, *E(spl)m8–MS2* transcription spanned a 25–30-minute window for each cluster (25.90 ± 4.9 min, mean ± SD), with an average duration per nucleus of circa 15 minutes (Figure 1E; S1C). During this phase, the apical area of the NB decreased until it fully delaminated (Figure 1C,F). This final stage of NB ingression was accompanied by an asynchronous loss in cluster transcription, which persisted in some cells for an additional 5–6 minutes (Figure 1F).

To investigate whether all cells within a cluster, including the NB, initially activate transcription, the clusters were backtracked to a stage several minutes prior to appearance of the first transcription foci. We then assessed transcriptional activity in the presumptive NB and the neighbouring cells (NCs) of each cluster (Figure 1C,E; Video S2). Based on previous analysis, we estimate that foci with 3 RNAs are above the detection limit^25^ yet no foci were ever visible in NBs either before or after transcription was initiated in NCs (Figure 1E). These results argue that there is a bias present within the proneural cluster that precedes the onset of *E(spl)m8* transcription and that shapes initiation of Notch signalling within clusters.

We next asked whether a second Notch-responsive gene, *E(spl)m7*, exhibited the same transcriptional dynamics, using a similar *E(spl)m7–MS2* line^25^. When aligned by transcription onset, *E(spl)m7–MS2* produced comparable transcription profile intensities and durations to *E(spl)m8–MS2* (32.23 ± 5.6 mins;Figure 1G, S1C; Video S1). Importantly, *E(spl)m7–MS2* was only transcriptionally active in NCs and was silent in the presumptive NB throughout the entire signalling window. This reinforces the conclusion that Notch signalling never exceeds the threshold needed for inducing target gene expression in the NB.

Altogether, these observations reveal a dynamic pattern of Notch-responsive transcription over the timescale of the fate decision being executed. The absence of detectable transcription of *E(spl)m8–MS2* and *E(spl)m7–MS2* in the presumptive NB argues that there is a bias in signalling from the outset. As the two genes had similar properties we focused on *E(spl)m8–MS2* for subsequent experiments.

### Presumptive NB is the signalling source for cells in direct contact

There are two models to explain the pattern of transcription observed. One is that the presumptive NB is the signalling source, activating the pathway in NCs in direct contact. The other is that NCs signal equally to each other through mutual interactions while the NB is blocked from responding.

To test experimentally which cells provide the signal, we carried out targeted laser ablations during the period when transcription first initiates (0–5 minutes), removing either the NB or an NC. Proneural clusters were identified using a reporter containing 3kb proximal *achaete* regulatory sequences upstream of Halo coding sequences (*ac-Halo*). This was combined with *E(spl)m8–MS2* and at the stage when transcription foci first appeared ablations were focused either on the NB or an adjacent NC. The cluster was then tracked to monitor the consequences.

Removal of the presumptive NB led to a rapid decline in *E(spl)m8–MS2* intensity within all adjacent NCs within 5–6 minutes, demonstrating that the NB provides the signal needed to sustain transcription in NCs. In contrast, ablating an individual NC did not diminish transcription in adjacent cells. Transcription persisted robustly in the adjacent NC and only declined after its regular duration (Figure 2A,B). This suggests that NCs do not act as significant sources of signal. Rather, they behave as recipients of NB-derived signalling.

**Figure 2.**
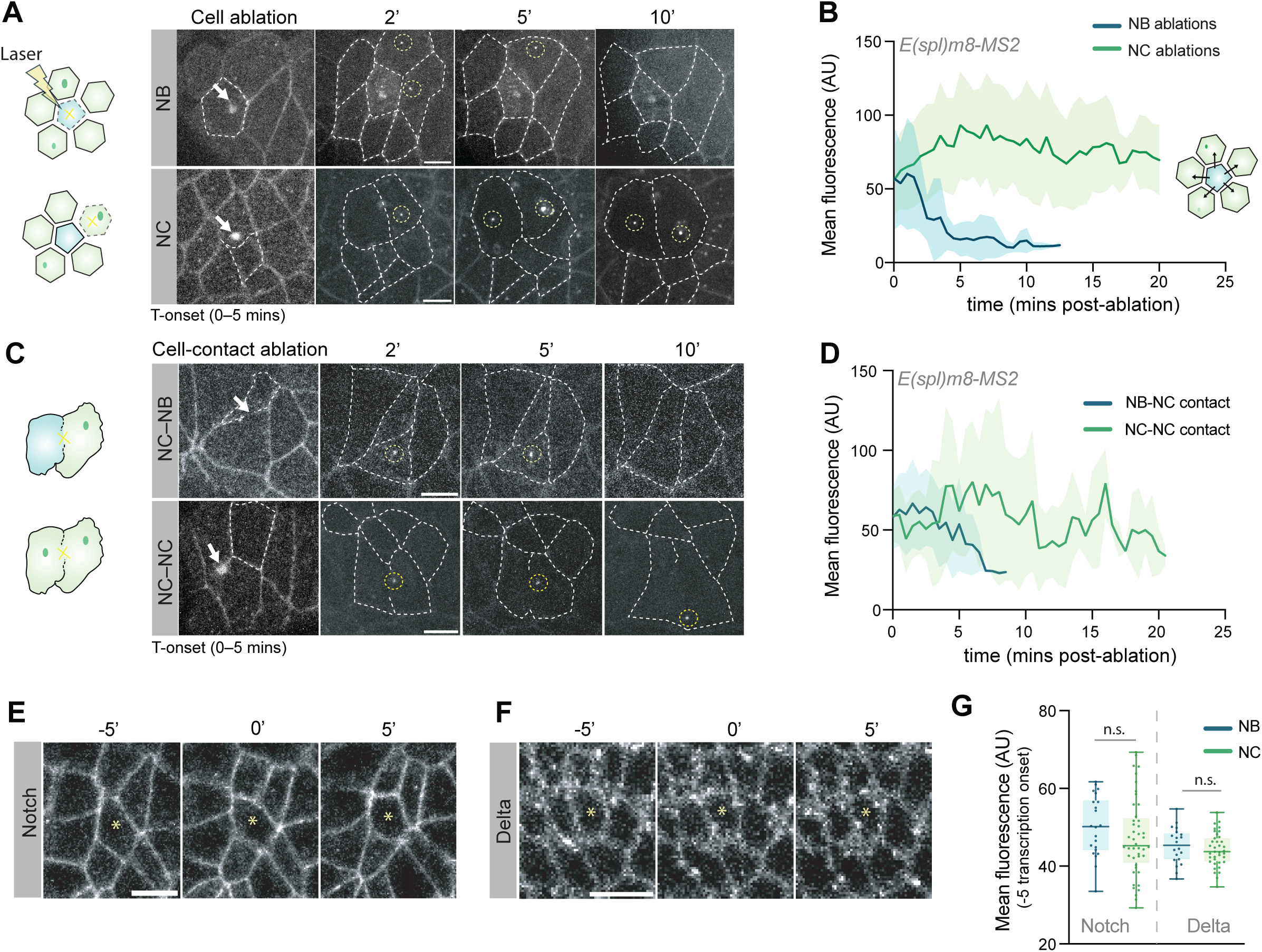
Presumptive Neuroblast (NB) is the primary signal-sending cell within proneural clusters. **(A)** Schematic summarising targeted single-cell laser ablations and representative images from each ablation type, executed immediately after T-onset (0–5 min). White arrows indicate ablated cells; dotted lines outline cell boundaries; yellow circles mark *E(spl)m8-MS2*/MCP transcription foci. NBs are outlined in the cell ablation panel. NB ablation leads to transcription decline in adjacent NC. Scale bars, 10 µm. **(B)** Mean *E(spl)m8–MS2/MCP* fluorescence traces over time following ablation of NB (blue) or NC (green). NB ablation results in loss of NC transcription, NC ablation does not. Mean transcription duration with NB ablated, 9 ± 2.08 mins (N= 16), and with NC ablated, 16 ± 3.56 mins (N= 26). Shaded regions indicate SD. **(C)** Schematic summarising cell-contact laser ablations and representative images from each ablation type, selectively removing either NB–NC interfaces (top) or NC–NC interfaces (bottom) immediately after T-onset (0–5 min). NB–NC contact ablation leads to loss of transcription in the affected NC. **(D)** Mean *E(spl)m8–MS2/MCP* fluorescence over time following NB–NC (blue) or NC–NC (green) contact ablations. NB–NC contact ablation results in rapid transcriptional decline. Mean transcription duration with NB ablated, 6 ± 1.7 mins, (N=10) and with adjacent NC ablated, 13 ± 3.9 mins (N= 10). Shaded regions indicate SD. **(E–F)** Representative images from time-lapse imaging of **(E)** Notch::Halo and **(F)** Delta::mScarlet in proneural clusters before (-5’) and after (5’) transcription initiates (0’). Notch or Delta levels in NB (yellow asterisk) are comparable with NCs. Scale bars, 10 µm. **(G)** Mean fluorescence intensities of Notch::Halo (left) and Delta::mScarlet (right) at membranes of NB (blue) versus NC (green) cells at t = -5 mins (pre) relative to transcription onset. Boxplot shows median and interquartile range, dots represent individual NC or NB measurements. No significant differences are detected at these stages. p = 0.2422 (Notch) and 0.4005 (Delta).

As Notch activation depends on direct ligand–receptor engagement at cell-contacts^18,30^, we next performed spatially targeted cell-contact ablations to directly perturb contacts between the presumptive NB and an adjacent NC or between two NCs. NB–NC contact ablations led to a rapid decrease in *E(spl)m8–MS2* transcription intensities within ∼5 mins. In contrast, ablation of NC–NC interfaces had no detectable effect on *E(spl)m8–MS2* activity in the affected cells (Figure 2C,D; Video S3). Thus, NC-NC interactions do not appear to contribute to the timing or levels of Notch-responsive transcription.

Together, these observations establish that the NB is the primary source of signal activating Notch in NCs suggesting that productive signalling only occurs at substantial NB–NC interfaces and not at NC–NC interfaces. To investigate whether this correlates with a difference in ligand and/or receptor expression we examined the levels of endogenously tagged Notch::Halo and Delta::mScarlet. However, in neither case were the levels detectably different between NB and NCs during the pre-transcription phase or at transcription onset (Figure 2E–G). Thus, the transcriptional response can’t be attributed to differential ligand or receptor expression. This agrees with observations that NB selection can occur under conditions where Notch or Delta are supplied uniformly under exogenous transcriptional control^7^.

### A bias in apical size prefigures transcriptional onset within clusters

Previous analyses have shown that differences in cell size, contact length and/or contact durations can shape the signalling profile, with smaller cells tending to become the signal producing cells in some contexts^16^. We therefore asked whether there are morphological differences between cells in a proneural cluster that prefigure the bias in Notch dependent transcription.

First, all cells within clusters were segmented and tracked in real-time. Next, an integrated quantitative analysis was performed measuring apical area, contact lengths of cell interfaces, duration of cell-cell contacts and *E(spl)m8–MS2* transcriptional dynamics at single-cell resolution (Figure 3A–B). To address whether any differences precede transcription onset, we first focused on a time-window that spanned 2.5 minutes prior to transcription onset (Figure 3A), beginning just as mesoderm invagination completed.

**Figure 3.**
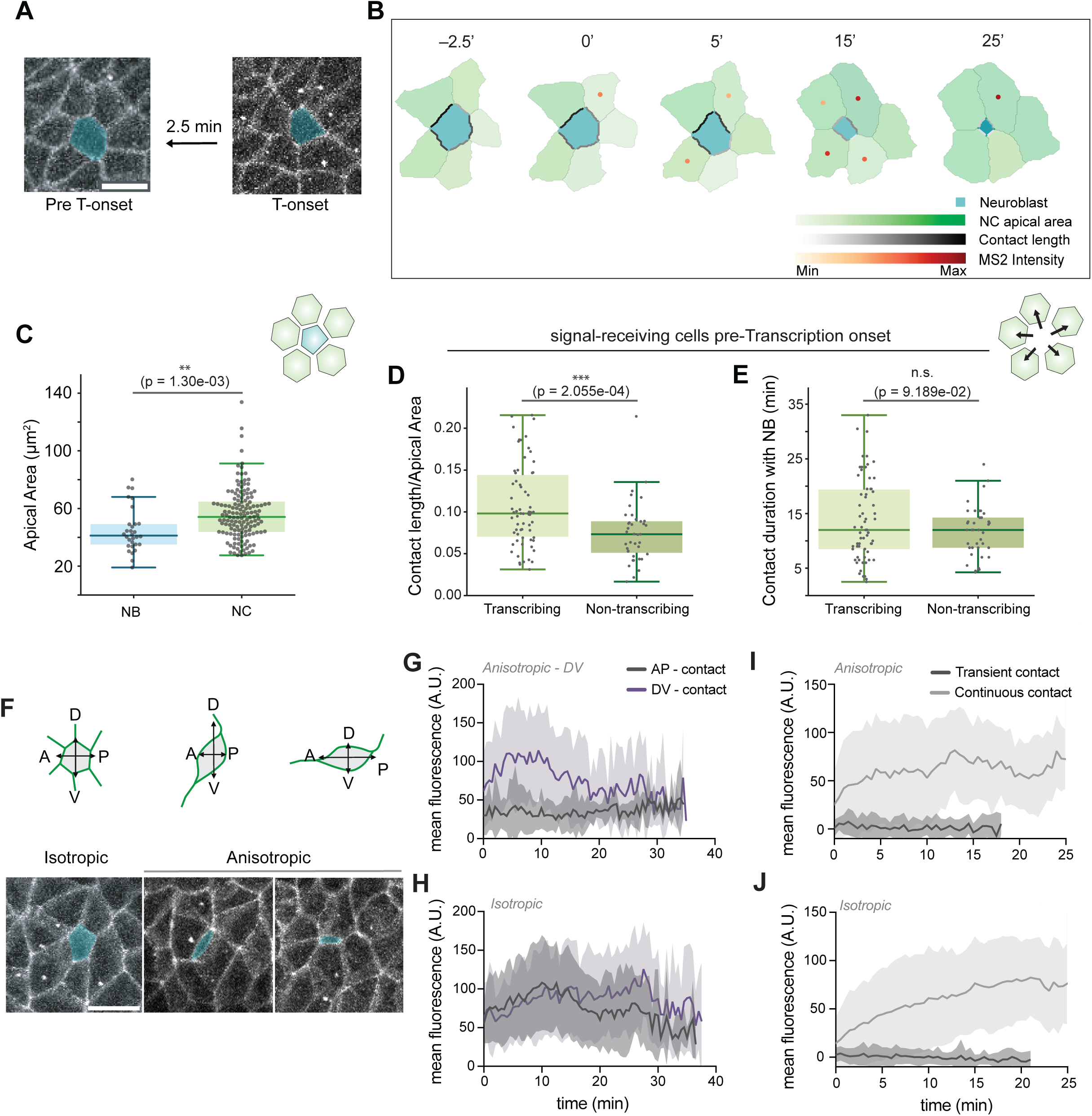
Quantifying apical areas and cell contacts within proneural clusters reveals a bias that pre-figures transcriptional outcome. **(A)** Images from a representative cluster before transcription onset (Pre T-onset) and at time of first detectable puncta of *E(spl)m8–MS2/MCP* transcription (T-onset). Spider-GFP marks cell membranes, presumptive NB is shaded blue (Scale bar, 10 μm). **(B)** Maps depicting quantified apical area (green scale), contact lengths (black scale) and transcriptional activity (MS2 intensity, red scale) over time (–2.5’, 0’, 5’, 15’, 25’ mins) for a proneural cluster, NB is shaded in blue. Legend indicates the colour scale for each parameter as labelled. **(C)** Quantification of mean apical areas for presumptive NBs (blue) versus non-NB neighbours (NCs, green) over a 2.5-minute window before transcription onset. NBs exhibit significantly smaller apical areas (44.764 ± 15.283 μm², N=30) compared to NCs (56.192 ± 17.873 μm², p = 7.334e-04, N=30 neuroblasts, 144 NCs). **(D)** Mean NB-NC contact lengths (normalised by NC areas) during a 2.5-minute window before transcription onset in neighbours that became transcribing vs non-transcribing (see methods). Transcribing NCs exhibit larger normalised contact length (0.107 ± 0.049) than non-transcribing (0.075 ± 0.036). N=66 transcribing, 39 non-transcribing. **(E)** Duration of NB-NC contacts before transcription onset in NCs that became transcribing vs non-transcribing. Durations are not significantly different (11.788 ± 4.56 and 13.78 ± 7.434 minutes pre-transcription). N=66 transcribing, 39 non-transcribing. **(F)** Schematic summarising different NB delamination geometries with representative images illustrating each type, NB highlighted in blue and Spider-GFP marks cell membranes. Isotropic NB delaminations, contact is maintained with ∼5–6 NCs forming a symmetric rosette-like cluster. Anisotropic NB delaminations, contact is maintained with fewer (∼2) NCs and NB acquires elongated shape aligned along DV or AP axis. Scale bar: 10 μm. **(G–H)** Mean *E(spl)m8–MS2/MCP* fluorescence over time from NCs in isotropic versus DV anisotropic clusters, plotted according to their AP (dark grey) or DV (purple) alignment. In anisotropic clusters, cells aligned to long axis of NB (DV, purple) have higher intensity. N=22 isotropic, 8 anisotropic clusters. **(I–J)** Mean *E(spl)m8–MS2/MCP* fluorescence over time from NCs in continuous contact with NB (light grey) versus those with transient contacts (dark grey) for isotropic and anisotropic clusters. TCs with transient contacts exhibit little or no transcription irrespective of delamination geometry. Isotropic, N=69 continuous, 40 transient-contact NCs; Anisotropic, N=15 continuous, 20 transient-contacts NCs.

Prior to *E(spl)m8–MS2* transcription initiation, the cells in each cluster appeared broadly equivalent in size, the overt visible decrease in the size of the NB became evident circa 5’ later as it commenced the contractions leading to its ingression. However, the quantitative analysis revealed a significant difference was present much earlier. The apical area of the presumptive NB was already smaller on average than that of other cells in the cluster even prior to the start of *E(spl)m8–MS2* transcription (Figure 3C). In situations where clusters contained two cells with smaller than average apical areas, the two usually became separated in time and the one remaining within the cluster went on to delaminate (Figure S2A). Thus, the bias in *E(spl)m8–MS2* transcription is preceded by a size differential, with smaller apical area prefiguring the cell that becomes signal sending and emerges as the single NB from the cluster.

To assess whether the cell contacts influence the probability of *E(spl)m8–MS2* transcription, we quantified the length and duration of the NB-interface for each NC in the cluster in the period prior to transcription onset. This revealed that the cells that initiated *E(spl)m8–MS2* transcription had a larger NB-NC interface prior to transcription onset than those that did not (Figure 3D) while the duration of those contacts did not differ significantly (Figure 3E).

We next asked whether NB interface contact length, contact duration, and NC apical area could discriminate which cells within the proneural cluster would initiate *E(spl)m8–MS2* transcription, indicative of Notch activation. Receiver operating characteristic (ROC) analysis was performed for each characteristic independently to descriptively quantify its discriminatory ability (Figure S2B). The length of the interface with the NB emerged as the strongest individual classifier (area under the ROC curve, AUC = 0.76), whereas contact duration showed only weak predictive ability (AUC = 0.55). NC apical area appeared non-predictive (AUC = 0.52∼0.5).

To further explore combinatorial effects from the different properties, we evaluated joint predictive performance using two supervised classification approaches (logistic regression and random forests), trained using repeated stratified 70/30 train–test splits to assess out-of-sample performance. Multivariate classifiers integrating all features achieved robust predictive performance (best model mean AUC= 0.749±0.067; Figure S2C,D), suggesting that these morphological characteristics relate to the probability that Notch signalling surpasses the threshold to activate transcription. Assessing the unique contribution of each predictor within the multivariate models using Permutation Feature Importance analysis confirmed that NB–NC interface contact length was the dominant driver of model performance, with contact duration making only a mild contribution (Figure S2E).

### Sustained contact with presumptive Neuroblast is required for transcriptional activity

Even if NB-NC contact duration is not a sufficient predictor for transcription initiation, a threshold duration may be required because most NCs that rapidly lost contact, due to rearrangements, failed to initiate transcription. This was most evident in clusters where the NB acquired an oval or lentil shape, indicative of a differential contraction of the cell interfaces. In these anisotropic clusters^23,31^ the long axis of the ingressing NB, which aligned with either the AP or DV axis of the embryo, retained contacts with two flanking cells (Figure 3F). In line with the ROC and multivariate classifier analyses, *E(spl)m8–MS2* was only robustly switched on in the cells in contact with the long axis of the NB (Figure 3G,H). In clusters where NB ingression was isotropic, contacting cells exhibited similar levels of transcription irrespective of their DV or AP positions (Figure 3I,J).

NCs in anisotropic clusters that were not aligned with the long axis of the NB rapidly lost contact, retaining at most a tricellular point contact at the NB interface. We next sorted the transcription profiles according to whether the cells had a transient or lasting contact with the NB. This confirmed that those with transient contact failed to initiate *E(spl)m8–MS2* transcription while those with sustained contact became transcriptionally active (Figure 3H). Similar behaviours were seen in isotropic clusters where the small subset of neighbouring cells that lost contact with the NB after early time points failed to initiate transcription (Figure 3J).

These results support the model that a threshold size/duration of contact is necessary to generate sufficient signalling to initiate transcription. Notably however, although cells that retained contact with the long axis of NBs in anisotropic clusters had longer interfaces than their isotropic counterparts, they did not achieve a higher mean intensity of *E(spl)m8–MS2* transcription (Figure G,H). This argues against a direct relationship between the length of the NB-interface and the levels of Notch-responsive transcription per se.

### Mechanical properties and delamination of Neuroblasts shapes the transcriptional response

Since Notch signalling is mechano-sensitive, one possible explanation for the role of cell shape and contacts in biasing signalling is that they modulate Notch activation via effects on junctional tension. Previous studies have shown that, at later stages when apical contraction of the NB is pronounced, NB-NC cell-cell contacts are under significantly higher tension than NC-NC contacts^23^. To establish whether there are mechanical differences even at earlier stages, we focussed on the time when *E(spl)m8–MS2* transcription was first detected in 1 cell per cluster. We performed targeted laser ablation on individual cell-cell interfaces and measured the initial recoil of tri-cellular junctions on both ends as an indication of tension along the cell-cell contact^32^ (Figure 4A). Comparing the recoil at NB–NC and NC-NC contacts we detected significant differences in tension from the earliest time measured. In all cases, NB-NC interfaces displayed higher recoil than NC–NC contacts (Figure 4B,C), indicating that there is already elevated tension at NB–NC interfaces when Notch dependent transcription is first detected.

**Figure 4.**
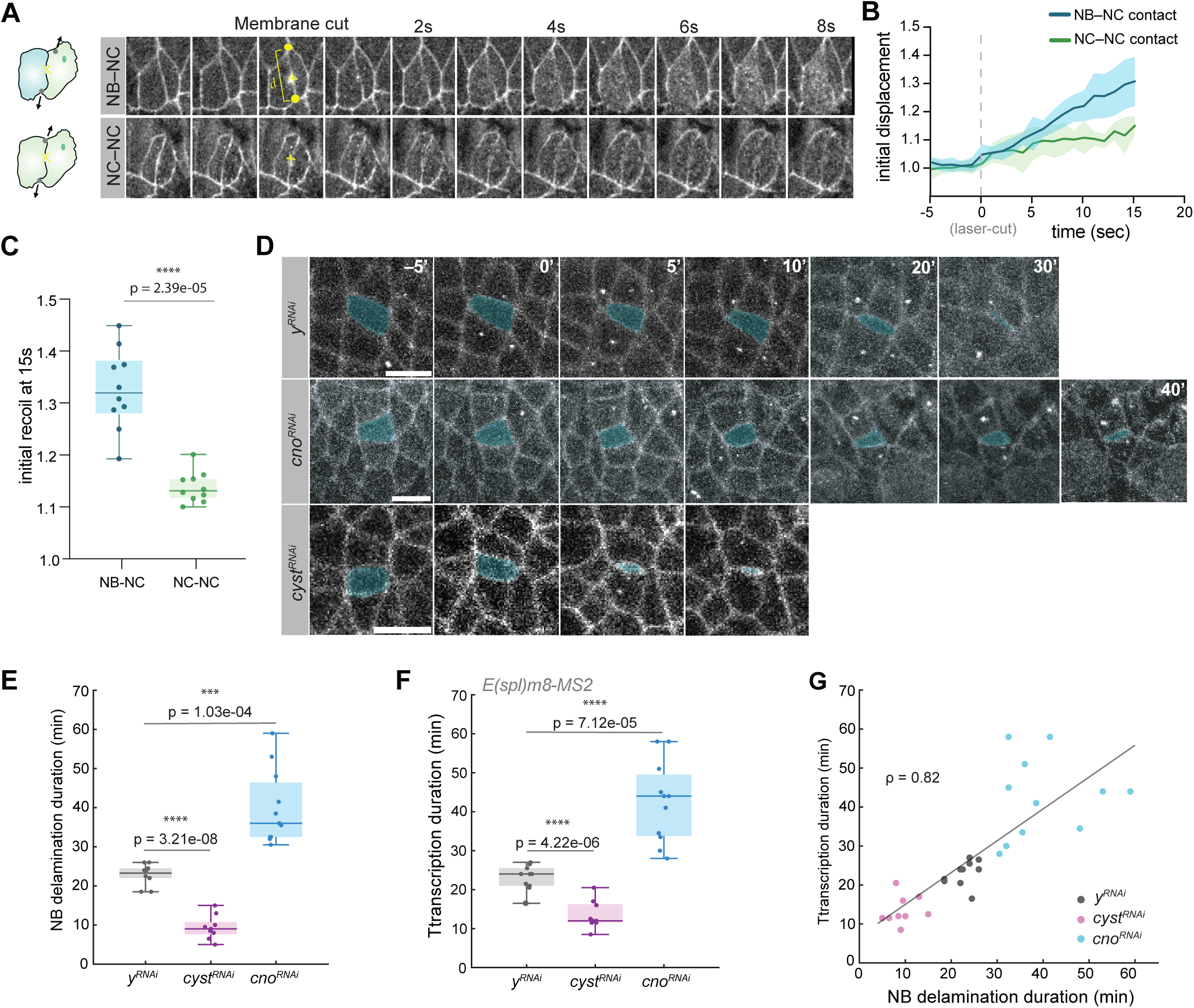
Role of junctional tension and ingression dynamics in modulating *E(spl)m8* transcription profiles. **(A)** Schematic summarising tension measurements (arrows) following cell-contact laser ablations and representative time-lapse frames (0–8 s post-laser cut) for each ablation type, selectively targeting either NB–NC interfaces (top) or NC–NC interfaces (bottom) immediately after T-onset (0–5 min). Recoil (*d*) of tricellular vertices (yellow dots) flanking each cut membrane was measured to assess junctional tension for NB–NC versus NC–NC contacts. **(B)** Quantification of normalized vertex displacement over time for NB–NC contacts (blue) and NC–NC contacts (green) upon ablation. Higher displacement values for NB–NC is indicative of higher junctional tensions. **(C)** Boxplot of initial recoil amplitudes at 15 sec post-ablation for NB–TC (blue) and TC-TC (green) interfaces. Dots represent individual measurements from contact recoil. NB–NC contacts exhibit higher recoil (1.3 ± 0.07) compared to NC-NC contacts (1.1±0.03) (p = 0.0041, Mann-Whitney U-test). N= 10. **(D)** Time-lapse images of *E(spl)m8–MS2* transcription in proneural clusters (NB in cyan) in *cno^RNAi^*, *cyst^RNAi^*and *y^RNAi^* (control) embryos. **(E)** Boxplots show delamination duration (time between transcription onset and NB disappearance from plane) across genotypes. Time taken for NB delamination is extended in *cno^RNAi^* embryos and decreased in *cyst^RNAi^* embryos. Boxplots N= 10, 9, 11 clusters for *y^RNAi^* (control), *cyst^RNAi^* and *cno^RNAi^* respectively. Mean delamination durations: 22.80 ± 2.78, 9.39 ± 3.07 and 39.91± 9.49 mins for *y^RNAi^* (control), *cyst^RNAi^* and *cno^RNAi^*respectively. **(F)** Quantification of cluster transcription duration (time between appearance of first MS2 spot and disappearance of last spot in each cluster) across genotypes. Transcription duration is extended in *cno^RNAi^* embryos and decreased in *cyst^RNAi^*embryos. N= 10, 9, 11 clusters for *y^RNAi^* (control), *cyst^RNAi^*and *cno^RNAi^* respectively. Mean transcription durations: 23.05 ± 3.20, 13.50 ± 3.64 and 42.45 ± 10.38 mins for *y^RNAi^*(control), *cyst^RNAi^* and *cno^RNAi^* respectively. **(G)** Correlation between NB delamination and cluster transcription duration (as quantified in E and F) across genotypes. Pearson correlation coefficient of 0.82 (p = 3.17e-08) indicates strong positive correlation.

To investigate further whether the mechanical properties contribute to the pattern of Notch-responsive transcription, we tested the consequences of perturbing factors known to regulate mechanical tension in the neuroepithelium. First, we depleted Canoe (cno), an adaptor protein that links actomyosin to cell junctions and is required for NB delamination to progress normally^23^. In *cno*-RNAi treated embryos, the prolonged period of NB delamination (39.91±9.49 min) was accompanied by an extended period of *E(spl)m8–MS2* transcription (42.45±10.38min; Figure 4D-E; Video S4). The strong correlation between the two suggests that they are related (Figure 4F,G). Similar effects were seen when embryos were injected with an inhibitor (BAY549) that targets Rho-kinase^33^, whose activity is required for Myosin-II contractility^34^. This treatment also prolonged the period of NB delamination (BAY-549= 34.06±6 min) and resulted in a similarly extended duration of *E(spl)m8–MS2* transcription (38.3±6 min vs 22.6±5 min controls; Figure S3A–C).

Lastly, we analysed the consequences from depleting the p114 Rho GTPase guanine nucleotide exchange factor (p114RhoGEF) Cysts, which activates Rho1 at adherens junctions and stabilizes junctional myosin. Its’ depletion reduces junctional myosin in the neuroepithelium, impacting on cell tensions, and leads to more rapid NB ingression ^35,36^. When Cysts levels were depleted, using *cyst-RNAi,* the accelerated NB ingression (9.39±3.07min; Figure 4D,E; Video S4)^35^ was accompanied by a reduced duration of *E(spl)m8–MS2* transcription (13.50±3.64min; Figure 4F,G). Together the data from *cno-RNAi* and *cyst-RNAi* confirm that there is a consistent relationship between the time taken for NB delamination and the duration of *E(spl)m8–MS2* transcription (Figure 4G).

### *E(spl)-m8* transcription correlates with morphological changes in Notch-active cells

Given the temporal correlation between *E(spl)m8* transcription and NB delamination, we explored the possibility that transcription levels were related to morphological changes in the NB and/or in the NCs. A cross-correlation analysis between *E(spl)m8–MS2* transcription profiles and the apical area of the associated NB over time (Figure 5A), revealed that NB apical size negatively correlated with the MS2 transcription intensity in NCs (Figure 5A). Thus, as the NB progressively constricted and lost apical area, the transcription levels in NCs increased. Since the anticorrelation decreased for negative time lags, the NB morphological changes do not precede the transcriptional activity but rather they are co-occurring.

**Figure 5.**
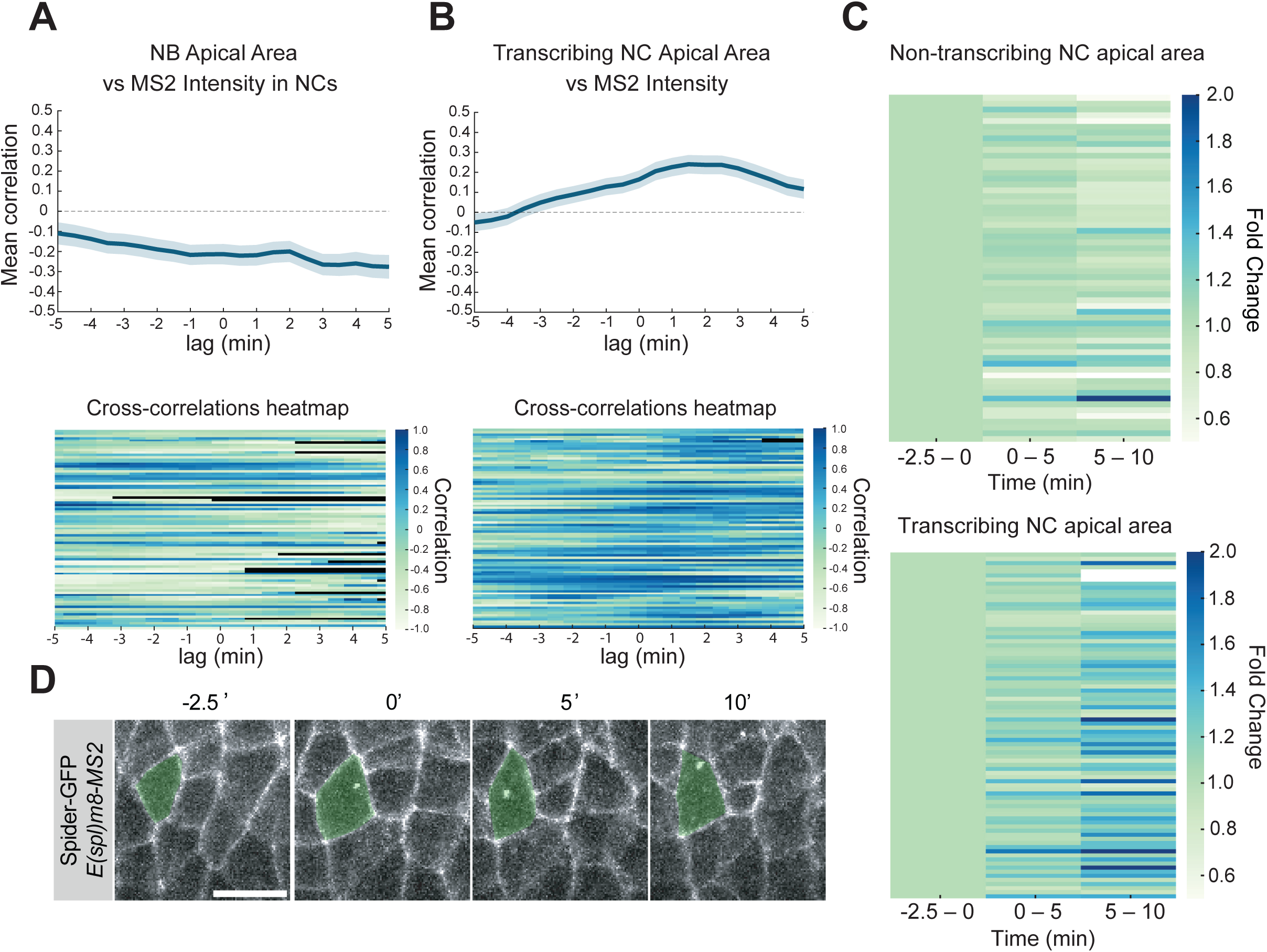
Relationship between *E(spl)m8* transcription levels and cell apical areas. **(A)** Cross-correlation analysis between NB apical areas and *E(spl)m8–MS2/MCP* transcription levels (fluorescence intensity) in NCs; upper, plot of mean correlation (over all NCs); bottom, heatmaps of individual cell cross-correlations. Negative correlation indicates the two are anti-correlated. **(B)** Cross-correlation analysis of the relationship between NC apical area and *E(spl)m8–MS2/MCP* transcription levels in each NC; upper, plot of mean correlation (over all NCs); bottom, heatmaps of individual cell cross-correlations. Positive correlation maximised at ∼2 mins indicates increase in NC area succeeds transcriptional increase in each transcribing NC. **(C)** Heat map indicating fold-change in apical area for transcribing and non-transcribing NCs averaged across two time-windows (0–5 min [1.10 +/- 0.15 transcribing, 0.99 +/- 0.16 non-transcribing], 5–10 min [1.28 +/- 0.33 transcribing, 0.95 +/- 0.26 non-transcribing] post transcription onset) in comparison to their average starting area at -2.5–0 min. Darker blue indicates large fold change. p-values: Transcribing: Window1 → Window2 p=4.10e-08, Window2 → Window3 p=1.46e-08. Non-transcribing: Window1 → Window2 p=6.01e-01, Window2 → Window3 p=3.98e-02 (paired t-tests). n=84 transcribing, 60 non-transcribing. **(D)** Representative images of apical-area dynamics of NC (green), before and after transcription onset. Transcribing NC apical areas increase during transcription.

In contrast, cross-correlation between the *E(spl)m8–MS2* intensities and apical area of the cognate transcribing cells revealed a positive correlation, which maximised at a positive time delay of ∼2min (Figure 5B). This implies that an increase in apical area follows the increase in mean *E(spl)m8–MS2* intensity in that cell. To investigate this further we quantified apical area trajectories of transcribing and non-transcribing neighbours in relation to the onset of transcription using three time-windows (2.5 mins prior to transcription onset, 0–5 mins and 5–10 mins after transcription onset; Figure 5C,D). Transcribing cells showed a marked increase in apical area during the 0–5 and 5-10 min windows following onset whereas their non-transcribing neighbours (cells that lose NB contact early or never reach a threshold contact length for activation) maintained relatively stable apical areas throughout the same period. In addition, there was a strong positive correlation between the apical area of a transcribing NC and its cumulative *E(spl)m8–MS2* level (area under MS2 curve, a proxy for mRNA numbers), for the majority of NCs (Figure S3D)

Thus *E(spl)m8–MS2* transcription in NCs reflects the progressive NB apical constriction and prefigures a significant apical expansion in the NCs themselves. The latter implies that Notch activation quantitatively impacts on the morphology of the cells within the cluster.

### Lateral inhibition model incorporating cell perimeter and tension-differences can replicate signalling properties

Our experimental analysis indicates that a bias in apical geometry and tension precedes transcription. The presumptive NB has a smaller apical perimeter and exhibits higher junctional tension than NCs. Furthermore, NCs transcribing *E(spl)m8* display a marked increase in apical perimeter that correlates with transcription levels.

These observations prompted us to test whether a modified lateral inhibition framework incorporating differences in apical cell perimeter and tension could account for the observed *E(spl)m8* expression pattern, including the absence of transcription in the presumptive NB even at the earliest stages.

Because Notch and Delta are predominantly localised to apical adherens junctions at this stage^26^, we implemented a two-dimensional model. It is based on a previously validated lateral inhibition model^15^ and has been modified to address the interplay between morphology, mechanics, and lateral inhibition. In the model, cells express Notch and Delta that can interact in trans to produce signalling (NICD) or in cis, causing cis-inhibition of receptors and ligands (Figure 6A). In contrast to the original model, we do not pre-suppose differential activator expression among cells. To account for contact-dependent signalling, we introduced perimeter-weighted cell–cell connectivity, which affects the function of Notch and Delta along cell boundaries (see Supplementary Information).

**Figure 6.**
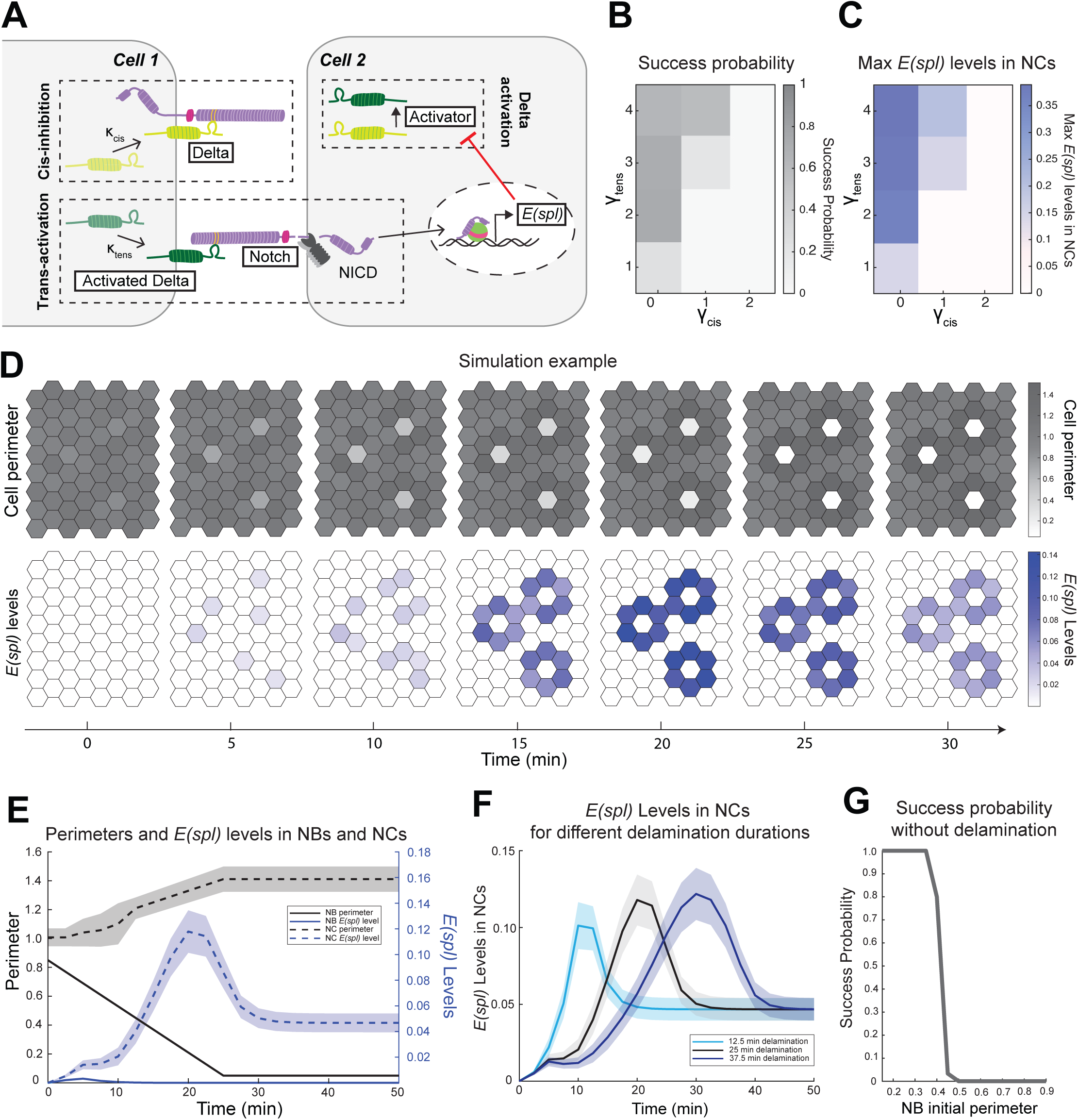
Incorporating morphological features into lateral inhibition model recapitulates in vivo *E(spl)* expression dynamics. **(A)** Schematic of lateral inhibition model, depicting cis-inhibition and transactivation of Notch by Delta. Trans-activation releases NICD, driving transcription of *E(spl)* genes leading to Delta activator inhibition. κ_cis_ scales rate of cis-inhibition, hypothesised to be inversely proportional to cell perimeter. κ_tens_ scales amount of activated Delta, hypothesised to be tension dependant and inversely proportional to cell perimeter. **(B)** Heatmap of simulation success probabilities, averaged over different *E(spl)* detection threshold values (see Supplementary). Success probabilities calculated across values of γ_cis_ and γ_tens_, the exponents that scale κ_cis_ and κ_tens_, respectively (simulations duration = 50mins). Darker shading indicates higher success probability. **(C)** Heatmap of maximum *E(spl)* levels in NCs from simulations, averaged over different *E(spl)* detection threshold values (see Supplementary). Maximum *E(spl)* levels are computed across values of γ_cis_ and γ_tens_. Darker shading indicates higher peak *E(spl)* expression. **(D)** Representative successful simulation of an 8×8 2D lattice, temporal changes in cell perimeters (top panel, grey scale) and *E(spl)* levels (bottom panel, blue scale), NC perimeter was set to increase once *E(spl)* expression surpassed a defined threshold. Parameters used: *E(spl)* detection limit = 0.015, initial perimeter for pre-NBs = 0.85, initial perimeter for all other cells = 1 (± 5% noise, see Supplementary), γ_cis_=1, γ_tens_=3. Delamination was enforced. **(E)** Quantification of perimeter (grey) and *E(spl)* levels (blue) over time for NBs (solid curves) and NCs (dashed curves) from the simulation shown in (D). NB delamination was set to occur over 25 min. *E(spl)* expression in NCs peaked during NB delamination. Only very low (below detection threshold) *E(spl)* expression observed in NBs. **(F)** Effect of NB delamination duration on NC *E(spl)* profiles, from representative simulations with 12.5 min (light blue), 25 min (grey; same as E), and 37.5 min (purple) delamination durations. Mean transcription durations over random seeds (time taken to reach E steady state): 21.20 ± 0.11min for 12.5min delamination, 33.02 ± 0.07min for 25 min delamination and 44.73 ± 0.08min for 37.5min delamination) **(G)** Simulation success probability for varying NB initial perimeter values in the absence of delamination. Parameters used for simulations: *E(spl)* detection limit = 0.012, γ_cis_=1, γ_tens_=3.

We next introduced two perimeter-dependent parameters. The first, a “tension” factor, κ_tens_, was based on the hypothesis that higher tension in smaller cells results in more active Delta at their boundaries, for example via effects on endocytosis. The second, a “cis-inhibition” factor κ_cis_, was based on the hypothesis that cis-inhibition scales inversely with cell perimeter, e.g. due to membrane crowding effects that limit availability of free Notch (Figure 6A; see Supplementary Information). These modifications allowed mechanical and geometric differences to influence signalling interactions directly.

To incorporate the observed changes in the presumptive NB as it undergoes delamination and the apical expansion of NCs that initiate *E(spl)* transcription, we imposed (i) a time-dependent decrease in NB perimeter and (ii) an increase in perimeter in cells whose *E(spl)* levels exceeded the detection threshold (see Supplementary Information). A simulation was scored as successful if the following criteria were met: (i) *E(spl)* expression occurred only in NCs, with at least one NC exceeding the detection threshold; and (ii) the ratio between maximal *E(spl)* levels in NCs and the detection threshold exceeded 10, consistent with calibration of our MS2 system^25^ (example in Figure 6D).

To explore how our assumptions and parameters affected lateral-inhibition patterns, we ran simulations for different values of γ_tens_ and γ_cis_, which are the exponents determining the perimeter dependency of of κ_tens_ and κ_cis_ respectively (see Materials & Methods and Supplementary Information). These revealed that the probability of successful simulations, with no expression in NBs, and maximal *E(spl)* levels in NCs (Figure 6B,C) increased with higher values of γ_tens_ (i.e. stronger dependence of Delta activity on perimeter and therefore tension). In contrast, increasing γ_cis_ (i.e. stronger dependence of cis-inhibition on perimeter), reduced both success probability and peak *E(spl)* levels in NCs. However, profiles obtained with γ_cis_>0 included a decrease of *E(spl)* expression following its peak during NB delamination similar to those observed. Under realistic parameter combinations, the model thus reproduced key features of the in vivo pattern: dynamic and asynchronous *E(spl)* accumulation in NCs and absence of expression in the presumptive NB (Figure 6D,E; Video S5).

We next examined the role of delamination dynamics, by varying the rate at which NB perimeters reduced. This changed the time required to achieve maximal *E(spl)* levels in NCs in the same direction, with prolonged delamination delaying peak of *E(spl)* expression and vice versa, resembling the shifts we observed in experiments (Figure 6F and 4G). In the absence of NB delamination, successful simulations were only obtained when the initial size differential was unrealistically large (NB:other ratio <0.5).

We then asked whether similar outcomes could arise from fully dynamic perimeter behaviour rather than imposed changes. To test this, we introduced an ordinary differential equation governing perimeter evolution as a function of intracellular *E(spl)* and activate Delta levels (see Methods and Supplementary information). We assessed whether the experimentally measured initial perimeter differential was sufficient to drive pattern formation under these conditions (Figure S4). Additional success criteria were imposed: (i) NB perimeter had to decrease below 0.3 by the end of the simulation, ensuring NB delamination was recapitulated and (ii) the mean perimeter of non-NB, non-NC cells had to remain above 0.6, as they had no substantial area loss in our experiments.

Within a 10 timepoint simulation window (∼25 min, see Supplementary Information), successful recapitulation of experimental behaviour was achieved for an initial perimeter ratio of ≤0.75 (Figure S4A), which closely matches experimentally measured values (Figure 3). Larger initial differentials further increased success probability. In this dynamic model, *E(spl)* levels in NCs reached higher amplitudes than in the imposed-perimeter model and activation coincided temporally with NB delamination and with reinforcement of NC perimeter, consistent with experimental observations (Figure S4B,C). However, at longer simulated times, additional cells began expressing *E(spl)* and extra cells underwent delamination (Figure S4D), suggesting that regulatory mechanisms, beyond those included in the present model, restrict the outcome at later temporal stages. Indeed, in vivo many of the cells undergo mitosis^27^.

As with classical explanations of Notch-mediated lateral inhibition, our model invokes a feedback mechanism that increases Delta activity in the signal sending cell relative to receiving cells. Although no quantitative difference in Delta::mScarlet distribution between NB and NCs was detected at transcription onset as discussed above (Figure 2E–G) a change in Delta enrichment appeared subsequently. Once *E(spl)m8–MS2* transcription levels in NCs increased and the period of definitive NB delamination was underway, Delta::mScarlet started to accumulate in large puncta within the NB (Figure S5). These observations are consistent with a model linking the apical cell-cell contacts and tension changes to effects on Delta activity.

Together the results show that lateral inhibition models based on an initial geometric bias can yield the correct profile of *E(spl)* expression when coupled with tension- and cis-inhibition-dependent effects. This supports our proposal that mechanical asymmetries create a bias in Notch signalling that accounts for its exclusion from the NB.

## Discussion

Many cell-fate decisions occur concurrently with morphological changes, and it is becoming increasingly evident that the latter plays an active role in the decision-making process. For example, there is mounting evidence that Notch signalling can be modulated by global and local mechanical cues ^21,37^ as we find here. Investigating the dynamic properties of cells undergoing Notch signalling during neurogenesis, by measuring *E(spl)m8* transcription in real-time, we discover a relationship between the cell morphology and their signalling outcome. Both our experimental results and mathematical modelling suggest that morphological and mechanical asymmetries help prime the system before transcriptional feedback engages to reinforce the differences and bring about the different cell fates.

In the context of neurogenesis, Notch mediated lateral inhibition operates in a landscape of cells thought to have equal potential. According to this model, they would initially all signal to one another, mutually repressing neural potential, until an inequality emerged and then became amplified through the transcriptional feedback loop^2–4^. Our data clearly demonstrate that aspects of this model are incorrect. There is an early bias between cells in a proneural cluster that precedes the onset of signalling, as detected by *E(spl)m8* transcription. One cell within a cluster never initiates transcription and our ablation experiments demonstrate that it adopts signal-producing status. This occurs without evident differences in ligand expression, consistent with previous studies demonstrating that neural precursor selection can occur normally when ligands are supplied uniformly^7^. Strikingly however, there were biases in the apical areas of the cells already at this “pre-signalling” stage: those cells with smaller apical areas became the signal-producing cell and delaminated as a neuroblast. Furthermore, neighbouring cells that had longer and more long-lasting interfaces had higher probability of *E(spl)m8* transcription and in turn increased in size more. A similar positive feedback loop between cell-cell contact duration and cell-fate specification has been observed for Nodal signalling emphasizing that such relationships are likely of general importance^38^.

The concept that variations in cell contacts are important in shaping Notch signalling has emerged in systems, such as the chick inner ear, where smaller cells are more likely to differentiate into hair cells^16^ and intestinal stem cells where cell-type differentiation correlates with contact area^39^. The distance that ligands and receptors diffuse in the cell membrane before they endocytose is potentially a key factor in this relationship. Tension heterogeneities modulate Notch activity in other contexts^21,22,37^, and we found that the contacts with the presumptive NB were at higher tension than those between neighbours. In addition, our observation that cell perimeters increased in proportion to the levels of *E(spl)* transcription argues that there is a positive reinforcement loop, whereby Notch activity increases apical areas and likely reduces tension in neighbouring cells. Together these mechanisms would reinforce the initial heterogeneities. Indeed, it’s been suggested that expansion of the apical domain in retinal epithelial cells augments Notch activation by recruiting activators or diluting the effective concentration of inhibitors^40^. Models incorporating tension-dependent effects on Delta-Notch binding have also been invoked in other contexts where they can recapitulate many forms of patterning^41^.

A key feature of neuroblast fate specification is its delamination, or ingression, a process which normally lasts for circa 20-25 minutes and relies on the tension anisotropy between NBs and neighbours^23,24^. The period of *E(spl)* gene transcription in each proneural cluster closely mirrored the delamination time, suggesting the two are coordinated. Furthermore, manipulations that perturb cytoskeletal components involved in regulating tension, such as mutations in *cyst* and *canoe*^23,35,36^, had concomitant effects on delamination times and *E(spl)m8* expression. This argues that the NB and its morphological transitions are important in orchestrating the levels and duration of Notch signalling.

To explore whether heterogeneities in the tension and lengths of cell-cell contacts could be sufficient to bring about the observed dynamics of *E(spl)m8* expression, we incorporated them into a model of lateral inhibition^15^, reasoning they’d impact on levels of receptors and active ligands available for trans signalling and on the likelihood of cis-inhibitory interactions occurring^13–15^. By initiating the model with the small differences between cells that we detected prior to signalling we could replicate key features of *E(spl)* expression. Notably, no transcription was detected in the presumptive NB and *E(spl)* transcription appeared asynchronously only in cells neighbouring the NB. Thus, subtle morphological differences can bias lateral inhibition and become further enhanced through Notch-transcription dependant cell perimeter increases in NCs. A similar feedback loop between local tension and Notch-mediated lateral inhibition is implicated in differentiation and patterning within the developing myocardium^22^ suggesting this interplay may of widespread relevance.

The absence of any *E(spl)m8* transcription in the presumptive neuroblast was unexpected. It implies that Notch signalling acts on a pre-existing bias that predisposes one cell to become the predominant source of ligand activity. In our analysis, we found that this corresponded to the cell with the smaller apical area. In a related study, Green and Schweisguth have made a similar observation tracking the fate of cells expressing proneural proteins. Both studies reveal therefore that, during neurogenesis and likely in many other contexts, Notch signalling acts within a landscape that is already primed and serves to re-enforce an established bias. The concept that such bias is manifest in morphological differences that become reinforced through signalling offers a powerful way to integrate signalling with tissue architectures.

## Acknowledgements

We are grateful to Yan Yan (Hong Kong University of Science & Technology) for generously sharing the *achaete-halo* flies with us and to Carmen Santa Cruz Mateos for Notch::Halo. We thank Kat Millen for help with generation of flystrains and Lisa-Maria Needham for her support of the microscope development. This work was supported by an Investigator Award (212207/Z/18) from Wellcome Trust to SJB, by a grant from Isaac Newton Trust to Lisa Maria-Needham and SJB, by a Wellcome Trust Technology Development Grant (212936/Z/18/Z) for the photoablation microscope to MOL, and by a PDN-Wolfson College PhD studentship to CR. Stocks obtained from the Bloomington Drosophila Stock Center (NIH P40OD018537) were used in this study.

## Author Contributions

Conceptualization: PR, CR, DS, SJB

Methodology: PR, CR, CZ, MOL, SK, DS, SJB

Software: CR, CZ

Validation: PR, CR,

Formal Analysis: PR, CR,

Investigation: PR, CZ

Resources: PR, CR, CZ, MOL

Visualization: PR, CR, SJB

Supervision: SJB, MOL, DS

Writing—original draft: PR, CR, SJB

Writing—review & editing: PR, CR, CZ, MOL, SK, DS, SJB

Project Administration: MOL, SJB

Funding acquisition: SJB

## Materials & Methods

### Experimental model

*Drosophila melanogaster*. Flies were grown and maintained on food consisting of glucose 76 g/l, cornmeal flour 69 g/l, yeast 15 g/l, agar 4.5 g/l, and methylparaben 2.5 ml/l. *Drosophila* stocks and crosses were raised and maintained at 25°C unless stated otherwise. Embryos were collected on apple juice agar plates with yeast paste.

### Fly strains and Genetics

Details of *Drosophila melanogaster* strains used in this study are listed in Table 1, including those obtained from the Bloomington *Drosophila* Stock Center.

**Table 1:**
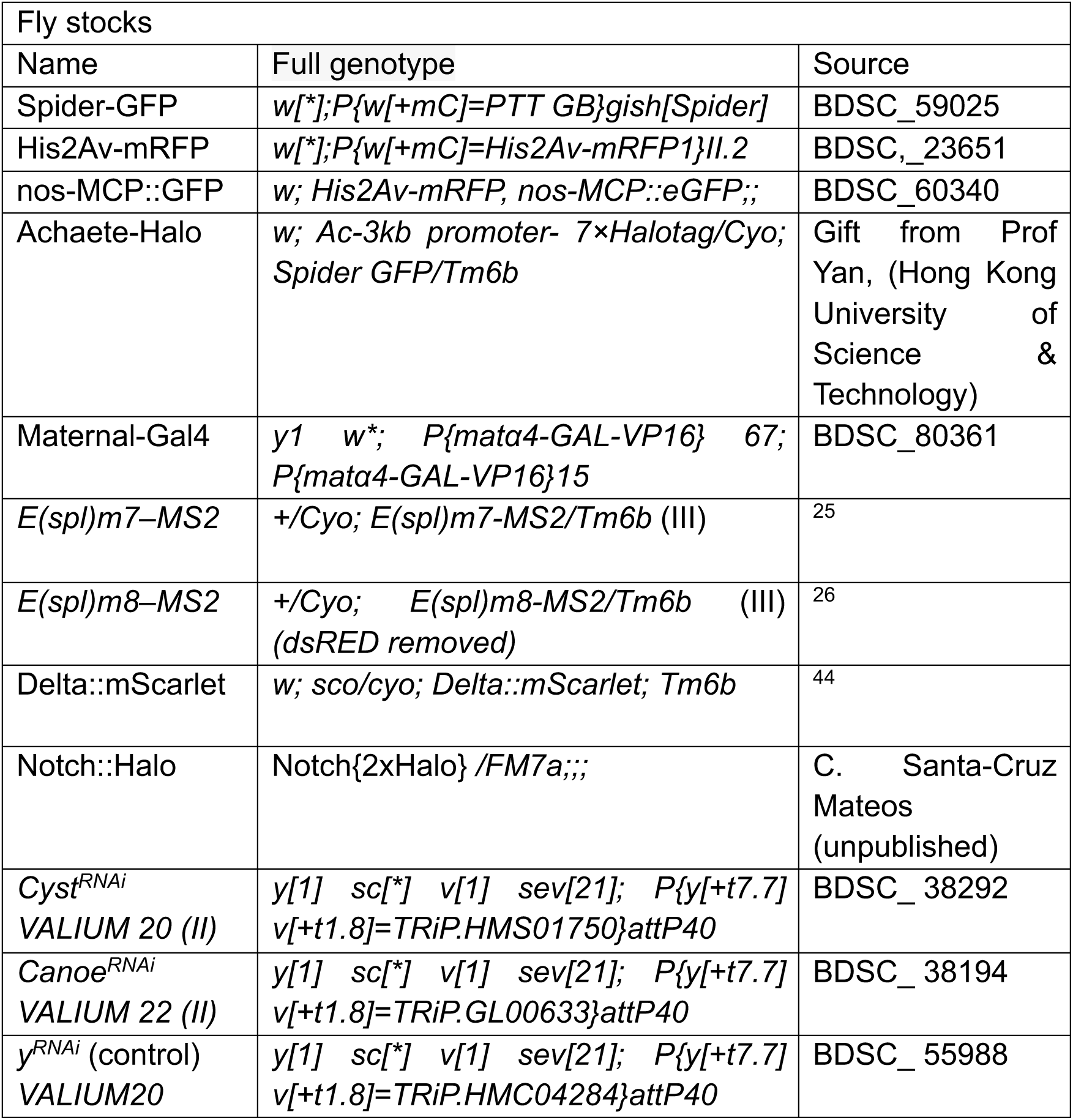
Drosophila strains.

To monitor transcription from the endogenous *E(spl)-C* locus, E(spl)*m8-MS2*^26^ *or E(spl)/m7-MS2*^25^ lines were crossed to *His2Av-RFP, nos-MCP::GFP; Spider-GFP* to generate embryos of the genotype: *His2Av-RFP*, *nos-MCP::GFP* (II); *E(spl)m8–MS2* / *Spider-GFP* (III). (Figure 1C).

To label proneural clusters in the neurectoderm, *Achaete-Halo (Ac-Halo); Spider-GFP* males were crossed to *His2Av-RFP, nos-MCP::GFP; E(spl)m8–MS2* females, generating embryos of genotype: *Achaete-Halo / nos-MCP::GFP (II); Spider-GFP / E(spl)m8–MS2 (III)*. (Figure S1A) *cyst^RNAi^, cno^RNAi^,* or *y^RNAi^* (control) were expressed maternally using the *mat-αTub-GAL4* driver (*mat-Gal4*). *mat-GAL4; Spider-GFP* females were crossed to *UAS-(cyst / cno / y)^RNAi^; nos-MCP::GFP, E(spl)m8–MS2* (recombined on the *III^rd^* chromosome) males to obtain embryos of the genotype: *mat-GAL4 / UAS-(cyst / cno / y)^RNAi^ (II); Spider-GFP / nos-MCP::GFP, E(spl)m8–MS2 (III).* (Figure 4D)

For experiments analysing Delta dynamics, males carrying *Delta::mScarlet* were crossed to *nos-MCP::GFP; E(spl)m8–MS2* females. (Figure S5 A)

All crosses were maintained under standard conditions, and embryos were collected and imaged at the indicated developmental stages.

### Mounting and live imaging of *Drosophila* embryos

0–2h, stage 6 embryos were collected on apple juice agar plates. Embryos were dechorionated using bleach and mounted onto 35-mm poly-D-lysine-coated glass bottom dishes (Mattek, P35GC-1.5-10-C). For all recordings (unless specified), embryos were oriented with their ventral/ventrolateral side facing the coverslip and submerged in 1x PBS.

Imaging was performed on a Leica SP8 confocal microscope (unless stated otherwise) using a 40x apochromatic, oil immersion objective (NA 1.3). Spider-GFP and nos-*MCP::GFP; E(spl)m8–MS2* detection were carried out using identical settings across experiments (except laser ablation experiments) and resolved based on their distinct localization within cells: A 40 mW 488-nm argon laser, and two hybrid GaAsP detectors, 8-bit, 600 Hz scanning speed, and a pinhole set to 4-Airy units. For imaging neurectodermal clusters, the following settings were used: 4% 488-nm laser power, 2x zoom, image size 512 x 512 pixel (0.43 μm/pixel), 1μm optical sections of 20–25 z-stacks, and a temporal resolution of 30s per volume. His2Av-RFP or HaloTag® TMR injected embryos were imaged with an additional 2%, 561-nm laser, image size 512 x 512 pixels resolution (0.43 μm/pixel).

### Embryo injections

HaloTag ligands: To image *Ac-Halo* and Notch::Halo, blastoderm stage embryos were collected and oriented ventral side facing the coverslip coated with mounting glue and were covered with halocarbon oil (Voltalef 10s). 0.25 mM of HaloTag® TMR (Promega G825A) or HaloTag® Ore-Green Promega G2801) ligands diluted in injection buffer (180 mM NaCl, 10 mM HEPES [pH 7.2], 5 mM KCl, 1 mM MgCl2) was injected into the perivitelline space of the embryos.

Rho-Kinase inhibitor: Embryos were injected with 2mM ROCK-inhibitor (BAY-549, Tocris Bioscience, TC-S 7001) or DMSO as control. To avoid interfering with ventral furrow formation, Rho-kinase inhibitor was injected into staged embryos immediately after the furrow had closed (early stage 7)

### Laser ablations

Embryos expressing *Ac-Halo*, *Spider-GFP* and *nos-MCP::GFP; E(spl)m8–MS2* were mounted on the coverslip to ablate single cell or precise cell-contacts. Gastrulation stage embryos were injected with HaloTag®TMR (Promega) 10mins before imaging. Proneural clusters within 0–5 min of transcription onset was located using TMR signal together with presence of Initial *E(spl)m8–MS2* spots.

### Optical arrangement of Photoablation unit

The imaging was performed on a Nikon Ti2 inverted-photomanipulation system. A confocal spinning disk system (CrestOptics X-light V3) is connected to the microscope. The confocal system is mainly composed of LDI-7 laser diode Illuminator (89 North Oxxius Inc), a lenslet array, a moveable spinning disk, two cameras (Photometrics Prime 95B), a 100X oil objective (Nikon CFI Plan-Apochromatic DM Lambda 100X Oil NA/1.45). The custom developed photoablation unit is composed of a femtosecond laser (Spectra-physics Mai Tai Deep Sea Ti:Sapphire HP), a polarizing beamsplitter, a Pockels cell (Lambda M350-105-BK-02), a beam expander, a spatial light modulator (Meadowlark 1920 x 1200). The ablation light is emitted from the Ti:Sapphire laser, polarized to plane waves, fast switch on and off by the Pockels cell and expanded to the desired beam size. The collimated beam is projected to the spatial light modulator for reshaping the beam profile to a tight spot, which is relayed to the image plane for ablation (<1 µm in diameter).

In a typical laser ablation experiment on *Drosophila* embryos, the laser wavelength was tuned to 900 nm to match 2-Photon absorption of GFP. Once the ablation site was precisely identified, the shutter was open for 15ms and closed for 20ms, 2 cycles for precise contact-ablations, 3 cycles for NB/NC ablations. The laser output power was set to result in 51 mW for precise contact ablations and 92 mW for NB/NC ablations on the image plane. The ablation process was controlled by a custom-developed LabVIEW (National Instruments) software, while the confocal system was controlled by MetaMorph (Molecular Devices) to continuously image the ablation and subsequent recoil process. Ablation was achieved by focusing the laser onto a single plane. For recoil measurements, single plane images were acquired immediately after ablation at the framerate of 1s/frame for 60s. For dynamic MS2-foci tracking, imaging was switched to acquire z-stacks with 30s/volume for 30 mins.

## Quantification and Statistical Analysis

### Cell segmentation and tracking

All image processing and quantitative analyses were performed using Fiji (ImageJ) together with MATLAB (Mathworks) or GraphPad-Prism. Cells were segmented and tracked and measured using the Tissue Analyzer integrated in Fiji^42^ Spider-GFP signal was used to segment cell boundaries and track apical areas, dynamic changes in cell–cell contact lengths and neighbour exchange within proneural clusters. Individual clusters were tracked starting at least 2.5 mins pre-transcription to 10 mins after apical NB delamination. NB and NCs were identified based on delaminating cells and presence of *E(spl)m8–MS2* dependent transcription and were backtracked to measure apical area, contact length and contact durations which were then plotted using a custom MATLAB script.

### Detection and quantification of transcription foci

Nascent transcription spots from *E(spl)m8–MS2 MCP::GFP* and *E(spl)m7–MS2 MCP::GFP* were tracked using the TrackMate plugin integrated in Fiji^43^. A LoG detector was applied with parameters optimised for the MS2/MCP::GFP foci size (normalized to the ROI diameter) and signal-to-noise characteristics and the mean-intensities of *MCP:GFP* spots were collected. Only transcription foci persisting for ≥ 2 consecutive frames were considered as transcription events. Final MS2/MCP::GFP intensities represent background-subtracted mean fluorescence values in clusters aligned to time 0 (i.e., detection of the first transcription foci to the appearance of last within a cluster).

### Combined analysis

Cell perimeter, contact-length, and MS2/MCP::GFP spot tracking obtained from TrackMate were combined using a custom MATLAB script to enable cluster-by-cluster analysis. This allowed simultaneous quantification of cell apical area, NB–interface contact length, and transcription levels over time.

For apical area comparisons shown in Figure 3C, the mean apical area of NBs and NCs was calculated over a 2.5 min window preceding cluster transcription onset. Onset was defined as the time when the first MS2/MCP::GFP puncta appeared within each cluster.

Contact length comparisons shown in Figure 3D were calculated as mean NB–NC contact lengths (normalised by NC area at each timepoint) during the 2.5 min window preceding the transcription start point. For transcribing NCs, this point corresponded to the time when the MS2/MCP::GFP puncta first appeared in that cell. For non-transcribing NCs, a pseudo-transcription start point was defined as the median transcription start time of the cluster.

Durations of NB–NC contacts shown in Figure 3E were measured from the time when the NC and NB first came into contact until the transcription start point of each transcribing NC, or the pseudo-transcription start point for non-transcribing NCs.

### Measurement of Notch and Delta intensities

Notch::Halo (Oregon Green ligand) and Delta::mScarlet intensities were measured in early-clusters 10 minutes prior to the continuous decrease in the apical-area of the NB. Measurements were done either at NB-NC contacts or NC-NC contact and normalized by subtracting background.

Embryos expressing Delta::mScarlet and *nos-MCP::GFP*; *E(spl)m8–MS2* were used to measure Delta dynamics within the clusters overtime. NBs or NCs were identified by the appearance of MS2/MCP::GFP puncta. Intensities from Delta vesicles localized in the cytoplasmic region of either NB or NC were measured and quantified excluding the signal from contacts-interfaces/membrane. Absolute intensity values were then normalized by subtracting the background.

### Recoil measurements

Junctional tension was measured as a proxy of the initial recoil dynamics following targeted laser ablation of individual cell-cell contacts. For each ablation event, continuous displacement of tricellular nodes at both ends of the junction were tracked using the line tool in Fiji. Displacement between each node was measured relative to the initial, pre-ablation distance between the nodes. Displacement trajectories were plotted for the first 15 sec following ablation and across multiple clusters to capture the instantaneous response at ablated contacts. Initial recoil at 15 sec was measured as a derivative of displacement over time.

### Cross-correlation analysis

Cross-correlation analyses (Figure 5A,B) were carried out using a custom MATLAB script, applying both negative (shifting MS2 curves backward in time) and positive (shifting MS2 curves forward in time) time lags.

For each time lag, the correlation between each shifted MS2 curve and its corresponding NB apical area (Figure 5A) or transcribing NC apical area (Figure 5B) was calculated (heatmaps in the bottom panels), and the average correlation across all tracks was then computed (curves in the top panels).

### Area fold-change analysis

Fold-change apical area analysis (Figure 5C) was calculated for transcribing and non-transcribing NCs across two time-windows, 0–5 min and 5–10 min after transcription onset, relative to their mean starting area during the −2.5–0 min window.

For non-transcribing NCs, the analysis was performed relative to cluster onset (the first time an MS2 spot appeared in the cluster) and continued for as long as the cluster persisted, irrespective of the NC’s contact with the NB.

For transcribing NCs, the analysis was performed relative to the transcription start point in that NC and continued for as long as transcription persisted in that cell, irrespective of NB contact.

### Modelling

We adapted the lateral inhibition model described in Troost *et al* ^15^ to investigate how cell geometry and mechanical properties influence signalling dynamics. In contrast to the original framework, we removed the requirement for a spatial activator gradient and instead implemented a perimeter-weighted cell–cell connectivity matrix. This modification allows the strength of interactions between neighbouring cells to depend on the extent of their shared boundary, reflecting the function of Notch and Delta along cell membranes.

To incorporate potential mechanical effects on signalling, we introduced two additional parameters which scale inversely with cell perimeter. The first parameter, κ_tens_, represents a “tension” factor that scales the amount of activated Delta available for trans-activation of neighbouring Notch receptors. The second parameter, κ_cis_, is a “cis-inhibition” scaling factor (Figure 6A; see Supplementary Information). These modifications allow cell geometry to modulate signalling interactions directly within the model.

The full mathematical formulation of the model is provided in Supplementary Information. This includes the reaction scheme corresponding to the processes illustrated in Figure 6A, the derivation of the governing ordinary differential equations, and the quasi steady-state assumptions used to simplify intermediate complexes. The Supplementary Material also describes the simulation framework, initial conditions, and parameter values used (see Supplementary Information and Table S2).

For analysis shown in Figure 6, the final set of non-dimensionalised equations used is as follows:

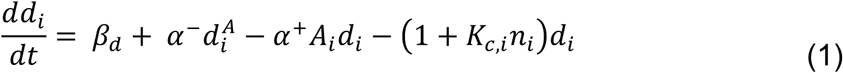

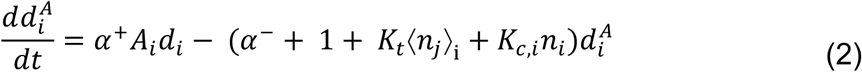

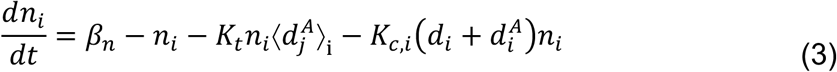

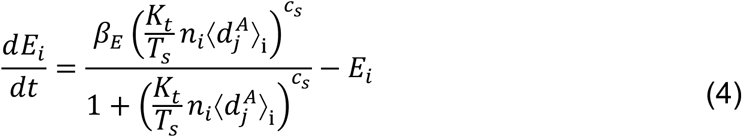

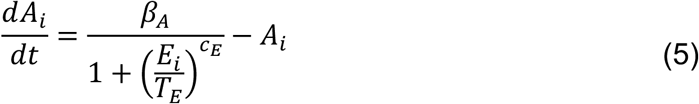

where 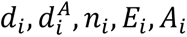, are levels of Delta, activated Delta, Notch, *E(spl)* and activator in cell i respectively,

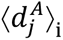 and 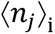 are sum of activated Delta and Notch, respectively, from cells j on the boundaries with cell i,

*β_d_, β_n_, β_E_, β_A_* are expression rates of Delta, Notch, *E(spl)* and activator respectively, *α* and *α*^-^ are the rates of Delta activation by Activator and de-activation respectively,

*K_c,i_* is the rate of cis-inhibition for cell i,

*K_t_* is the rate of trans-activation,

*T_s_* and *T_E_* are the Hill half occupation levels associated with Notch promoting *E(spl)* expression and *E(spl)* inhibiting activator expression respectively, while *c_S_*. and *c_E_*, are the respective Hill coefficients.

The following perimeter rule was set to account for NB delamination and transcribing NC apical area reinforcement:

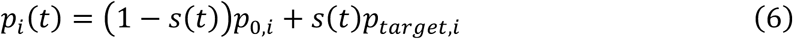

where,

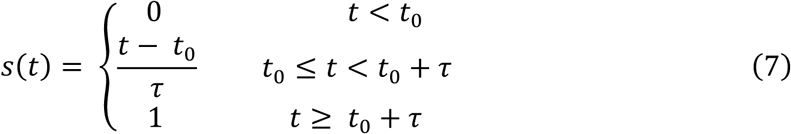

with *τ* the delamination duration and

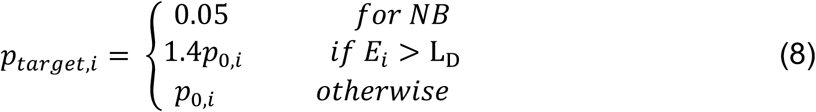

where *p_i_* is the perimeter of cell i (with *p_0_*_,*i*_ as the initial perimeter),

*τ* is the delamination duration and *t*_0_ the timepoint when this rule is triggered and L_D_ is *E(spl)* detection limit.

For the dynamic perimeter model (Figure S4), ODEs (1) – (5) were used together with a perimeter ode:

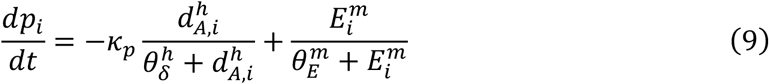

where *κ_p_* is the ratio of Activated Delta and *E(spl)* Hill function prefactors respectively,

*θ_δ_* and *θ_E_* are the Hill activation thresholds associated with Delta activity inhibiting and *E(spl)* promoting perimeter respectively, while *h* and *m* and the respective Hill coefficients.

The equations are solved numerically in MATLAB using a standard ODE solver.

### Descriptive statistics and statistical tests

Data are expressed as the mean ± SD, and error bars in graphs represent SD. For all boxplots, the line across the box represents the median, the top and bottom edges correspond to the upper and lower quartiles, respectively, and whiskers extend to 1.5× the interquartile range. Dots represent individual measurements.

For statistical tests involving two samples, two sample t tests (if samples were normal) and Mann-Whitney U tests (if samples were not normal) were performed. Normality of the samples was assessed using Q-Q plots and Shapiro-Wilk tests. Where two samples were compared, equality of variance was also assessed with Bartlett’s test (if samples were normal) and Levene’s test (if samples were not normal). In all cases significance was presented as follows: * p < 0.05, ** p < 0.01, *** p < 0.001, **** p < 0.0001.

## Data and code availability

All code used for the above-mentioned analyses and modelling can be found at: https://github.com/crou607/neurogenesis_analysis.git

## Supplemental information

Supplementary Legends and Figures S1–S5

Supplementary Information: **Additional details regarding computational model**

Supplementary tables: Table S2. Simulation framework, initial conditions, and parameter values. Related to Figure 6

Supplementary videos:

Video S1. *E(spl)m8–MS2* and *E(spl)m7–MS2* transcription dynamics in ventral ectoderm. Related to Figure 1, Figure S1C.

Video S2. Transcriptional activity within a cluster in relation to NB delamination. Related to Figure 1, Figure S1.

Video S3. Effects of Ablations on transcription activity. Related to Figure 2D F.

Video S4. Effects of RNAi depletions. Related to Figure 4D–G.

Video S5. Results from simulation Related to Figure 6.

