## Supplementary Information for "Notch mediated lateral inhibition is shaped by morphological differences to reinforce bias toward signal-sending or receiving roles"

**Document S1**

#### Supplementary Figures

Figure S1. Spatiotemporal coordination of cluster transcription. (Related to Figure 1)

Figure S2. Contact length predicts transcription within proneural clusters. (Related to Figure 3)

Figure S3. Delaying NB delamination via perturbing Myosin II based apical contractions prolongs transcription. (Related to Figure 4)

Figure S4: Dynamic perimeter model reproduces many features of *E(spl)* transcription. (Related to Figure 6)

Figure S5. Changes in Delta enrichment during transcriptional progression. (Related to Figure 6)

#### Supplementary Information for computer modelling

Table S1: Initial values and parameters for model

**Figure S1. Spatiotemporal coordination of cluster transcription. (Related to Figure 1)**

**(A)** Time-lapse images of embryo expressing *Achaete-Halo* reporter to mark a subset of proneural clusters (bright magenta, cytoplasmic) together with *Spider-GFP* (green) to visualise cell membrane, *E(spl)m8–MS2* transcription (green foci) and nuclear *H2Av-RFP* (dark magenta). *Achaete-Halo* can first be detected as ventral furrow completes closure, labelled t=0, and becomes stronger over time. Scale bar, 20 µm.

**(B)** Onset of *E(spl)m8–MS2* occurs asynchronously within proneural clusters. Heatmap in which each horizontal row corresponds to *E(spl)m8–MS2* fluorescence intensity in a single nucleus, aligned to the transcription onset within each cluster, N= 30 clusters.

**(C)** Transcription durations of individual *E(spl)m7–MS2* and *E(spl)m8–MS2* traces (all traces aligned to their onset). Boxplot shows median and interquartile range, dots represent individual measurements corresponding to a transcriptionally active NC. Both genes exhibit comparable on-duration of: m8 (16 ± 7 min) and m7 (21 ± 8 min).

### Figure S1

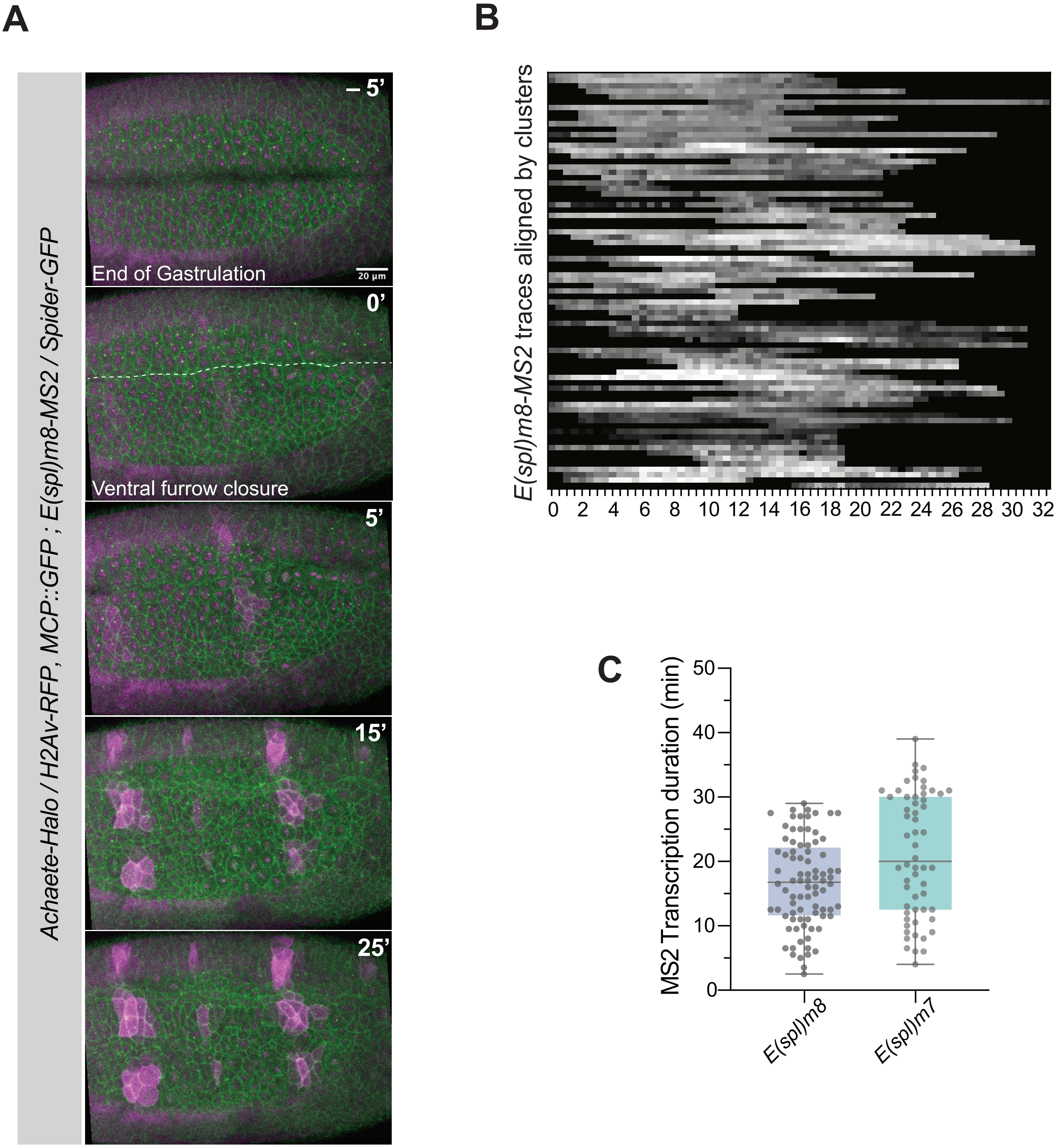

A
B
C
Achaete-Halo / H2Av-RFP, MCP::GFP ; E(spl)m8-MS2 / Spider-GFP
– 5'
End of Gastrulation
20 µm
0'
Ventral furrow closure
5'
15'
25'
E(spl)m8-MS2 traces aligned by clusters
0 2 4 6 8 10 12 14 16 18 20 22 24 26 28 30 32
250
200
150
100
50
MS2 Transcription duration (min)
50
40
30
20
10
0
E(spl)m8
E(spl)m7

**Figure S2. Contact length predicts transcription within proneural clusters. (Related to Figure 3)**

**(A)** Representative time-lapse images of two smaller cells (blue) initially in contact during pre-transcription stages that then undergo dynamic contact-rearrangements over time to participate in adjacent clusters and delaminate separately.

**(B)** Receiver operating characteristic (ROC) analysis performed independently for NB–interface contact-length, contact-duration, and NC apical area to quantify their ability to discriminate transcribing from non-transcribing cells. The area under each ROC curve (AUC) was used as a threshold-independent measure of performance, NB-interface length (dark blue curve) emerged as strongest individual classifier (AUC= 0.76), contact-duration (light-blue curve) showed weak predictive ability (AUC = 0.55). NC apical area (grey-curve) appeared non-predictive (AUC= 0.52).

**(C)** Joint predictive performance was quantified using two supervised classification approaches, Logistic Regression and Random Forests. Models were trained using repeated stratified 70/30 train–test splits (N= 20). Performance was assessed using ROC-AUC values calculated on held-out test sets. 0.749 ± 0.067 for Logistic Regression, 0.711 ± 0.078 for Random Forest.

**(D)** Logistic regression PCA decision boundary shown for a representative test set. Green and blue regions denote areas predicted by the model as non-transcribing and transcribing, respectively. Test set data are overlaid (green squares, non-transcribing cells; blue squares, transcribing cells) and the majority are correctly classified.

**(E)** Permutation feature importance computed on held-out test sets for logistic regression and random forest models. For each feature, values were randomly permuted and the resulting decrease in model performance (ROC-AUC) was measured. Larger decreases indicate greater predictive contribution. Results identify NB–interface contact length as the dominant driver of model performance. Logistic Regression: Contact length 0.243 ± 0.052, Contact duration: 0.032 ± 0.029, Apical area -0.016 ± 0.025. Random Forest: Contact length 0.143 ± 0.048, contact duration: 0.044 ± 0.035, apical area -0.014 ± 0.027.

### Figure S2

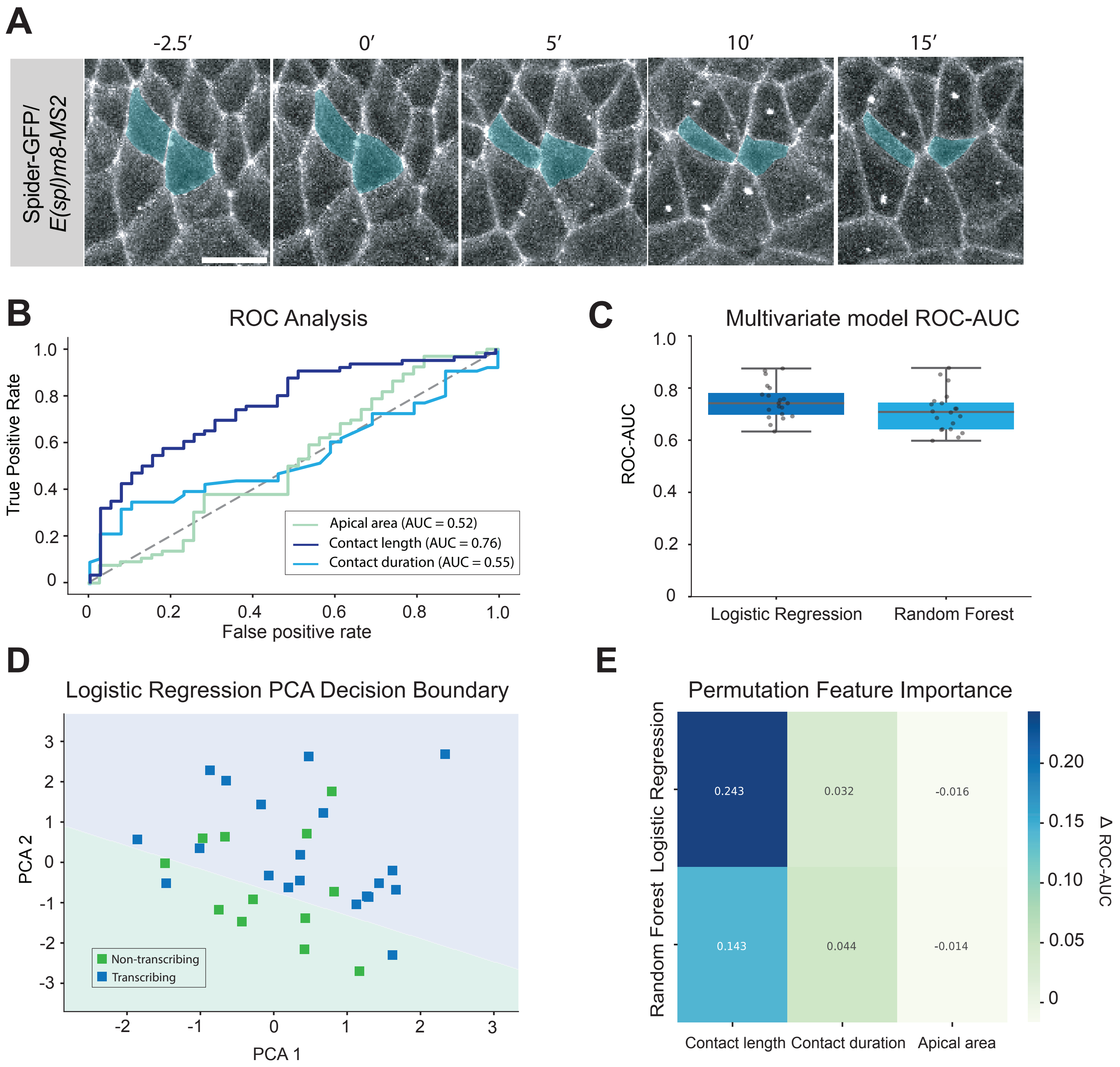
A
-2.5′
0′
5′
10′
15′
Spider-GFP/
E(spl)m8-MS2
B
ROC Analysis
True Positive Rate
False positive rate
Apical area (AUC = 0.52)
Contact length (AUC = 0.76)
Contact duration (AUC = 0.55)
C
Multivariate model ROC-AUC
ROC-AUC
Logistic Regression
Random Forest
D
Logistic Regression PCA Decision Boundary
PCA 2
PCA 1
Non-transcribing
Transcribing
E
Permutation Feature Importance
Logistic Regression
Random Forest
0.243
0.032
-0.016
0.143
0.044
-0.014
Contact length
Contact duration
Apical area
Δ ROC-AUC

**Figure S3. Delaying NB delamination via perturbing Myosin II based apical contractions prolongs transcription. (Related to Figure 4)**

**(A)** Time taken for NB to delaminate in DMSO (dark blue) and BAY549 ROCK-inhibitor (green) injected embryos, BAY-549 treatment significantly prolongs delamination duration N= 16 clusters for both DMSO and BAY549*.* Mean delamination durations: 25.25 ± 2.62, 34.25 ± 6.43 mins for DMSO and BAY549 respectively. p=4.64e-05 (two-tailed t-test). n=16 clusters for both DMSO and BAY549*.*

**(B)** Total duration of *E(spl)m8–MS2* transcription within proneural clusters in DMSO (grey) and BAY-549 (blue) treated embryos Cluster transcriptional duration is prolonged in BAY-549 treated embryos compared to the controls*.* Mean transcription durations: 20.28 ± 3.96, 44.66 ± 12.5 mins for DMSO and BAY549_respectively. p=7.12e-07 (two-tailed t-test). n=16 clusters for both DMSO and BAY549*.*

**(C)** Correlation between NB delamination and cluster transcription duration (as quantified in A and B) across DMSO and BAY549. Pearson correlation coefficient of 0.49 (p=0.0045) indicates positive correlation.

**(D)** Violin plot of correlations of NC Apical Areas vs area under MS2 curves (as a proxy for representing the cumulative mRNA numbers expressed in that NC). Mean ρ= 0.5632 ± 0.4072, indicates strong positive correlation for the majority of NCs.

### Figure S3

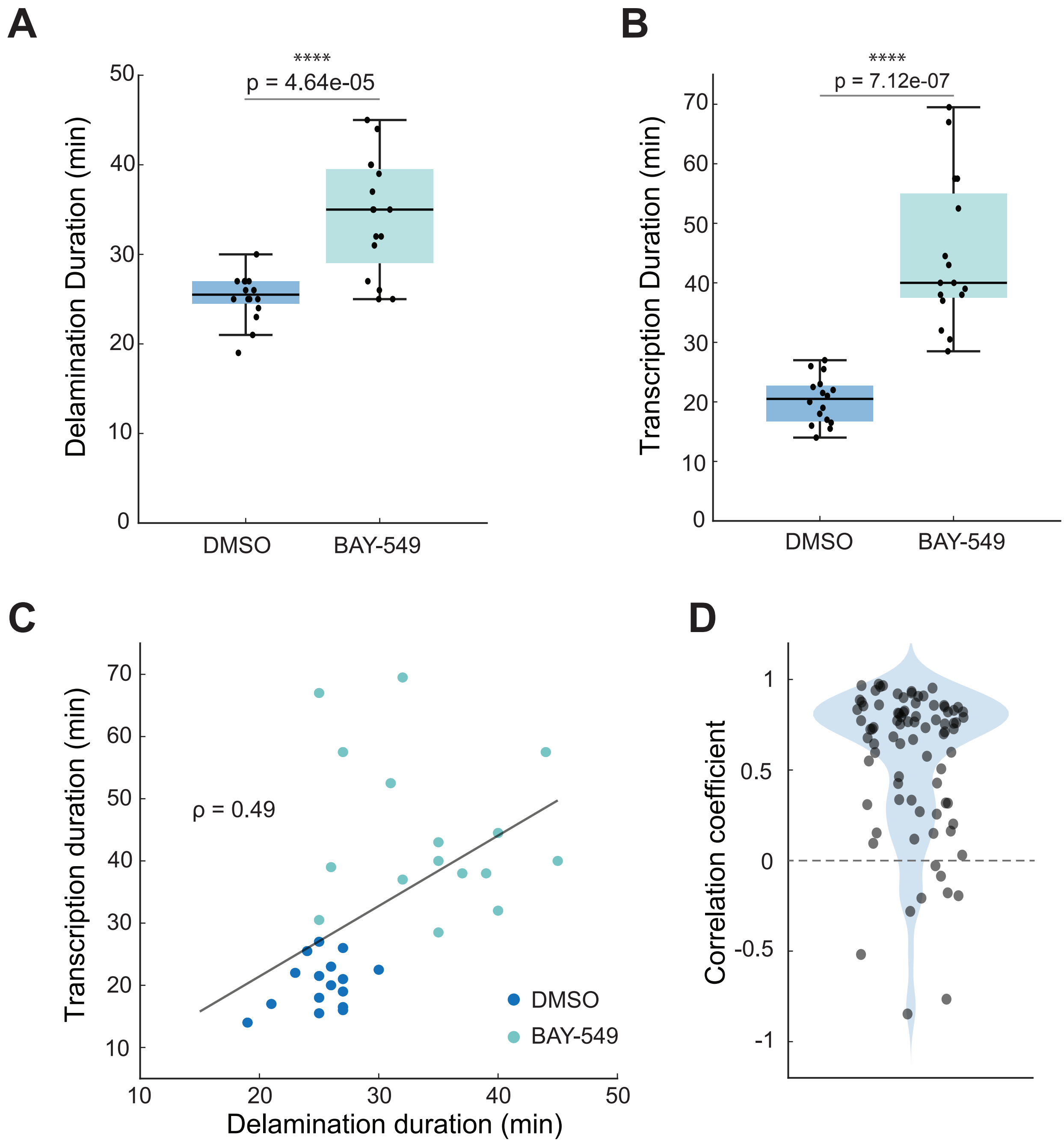
A
****
p = 4.64e-05
Delamination Duration (min)
0
10
20
30
40
50
DMSO
BAY-549
B
****
p = 7.12e-07
Transcription Duration (min)
0
10
20
30
40
50
60
70
DMSO
BAY-549
C
Transcription duration (min)
ρ = 0.49
DMSO
BAY-549
10
20
30
40
50
60
70
10
20
30
40
50
Delamination duration (min)
D
Correlation coefficient
1
0.5
0
-0.5
-1

**Figure S4: Dynamic perimeter model reproduces many features of E(spl) transcription. (Related to Figure 6)**

**(A)** Heatmap of simulation success probabilities using a dynamic perimeter model, in which cell perimeter depends on *E(spl)* and activated Delta levels within each cell. Probabilities are averaged over different *E(spl)* detection threshold values (see Supplementary Information) and are calculated across values of the initial NB perimeter and $\kappa_p$, the parameter that scales the effect of *E(spl)* and activated Delta on perimeter in each cell (simulations duration= 25mins). Darker colours indicate higher success probability (see main text and Supplementary for success criteria).

**(B)** Heatmap of maximum *E(spl)* levels in NCs from simulations with dynamic perimeter model, in which cell perimeters depend on *E(spl)* and activated Delta levels, averaged over different *E(spl)* detection threshold values (see Supplementary Information). Maximum *E(spl)* levels are computed across values of the initial NB perimeter and $\kappa_p$. Darker colours indicate higher peak *E(spl)* expression (see main text and Supplementary Information for success criteria).

**(C)** Quantification of perimeter (grey) and *E(spl)* levels (blue) over time for NBs (solid curves) and NCs (dashed curves) from representative simulation with the dynamic perimeter model. As observed experimentally, NBs delaminate over ~25 min, and NC perimeters increase with *E(spl)* expression. Once delamination completes, *E(spl)* levels in NCs decline, consistent with experimental measurements. Only very low (below detection limit) *E(spl)* expression is observed in NBs. Parameters used for the simulation: *E(spl)* detection limit= 0.025, initial perimeter for NBs=0.75, initial perimeter for all other cells= 1 (± 5% noise, see Supplementary methos), $\gamma_{cis}$=0, $\gamma_{tens}$=3. Perimeters are restricted between [0.05,1.4], if a cell perimeter reaches the boundary, dp/dt for that cell is set to 0.

**(F)** Successful simulation of an 8×8 2D lattice corresponding to (C), showing temporal changes in cell perimeters (top panel, grey scale) and *E(spl)* levels (bottom panel, blue scale). Delamination of 3 NBs is accompanied by asynchronous *E(spl)* expression in NCs at varying amplitudes. Experimental observations are recapitulated up to ~27.5 min, when delamination concludes, beyond which the simulation diverges. In vivo NCs would enter into mitosis, disrupting cell contacts and trans-signalling.

### Figure S4

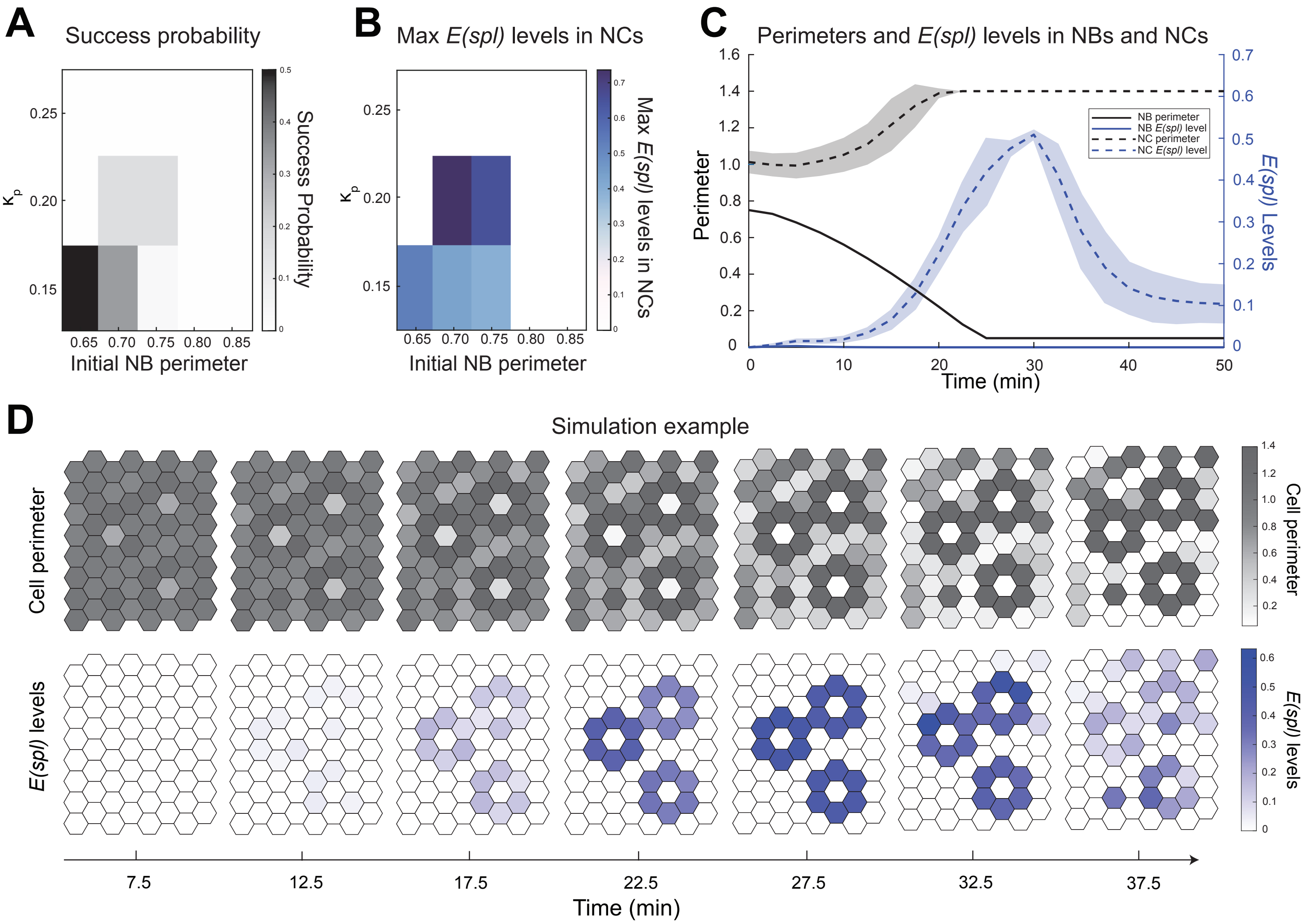

A
Success probability
0.25
0.20
0.15
Kp
0.65 0.70 0.75 0.80 0.85
Initial NB perimeter
Success Probability
B
Max E(spl) levels in NCs
0.25
0.20
0.15
Kp
0.65 0.70 0.75 0.80 0.85
Initial NB perimeter
Max E(spl) levels in NCs
C
Perimeters and E(spl) levels in NBs and NCs
Perimeter
E(spl) Levels
Time (min)
NB perimeter
NB E(spl) level
NC perimeter
NC E(spl) level
D
Simulation example
Cell perimeter
E(spl) levels
7.5
12.5
17.5
22.5
27.5
32.5
37.5
Time (min)

**Figure S5. Changes in Delta enrichment during transcriptional progression. (Related to Figure 6)**

**(A)** Representative time-lapse images of a proneural cluster expressing Delta::mScarlet (top row) and *E(spl)m8–MS2* transcriptional (middle row). Gradual enrichment of cytoplasmic Delta::mScarlet puncta occurs once transcription proceeds in NCs (Scale bar, 10 μm)

**(B)** Quantification of mean Delta fluorescence intensity over time in the NB (magenta) and neighbouring NCs (green), aligned to transcription onset (dashed line at 0'). Delta levels progressively increase in the NB post-onset of *E(spl)m8–MS2* transcription within the clusters, whereas NCs maintain relatively stable levels. Shaded regions indicate SD. (N=11 clusters).

**Figure S5**

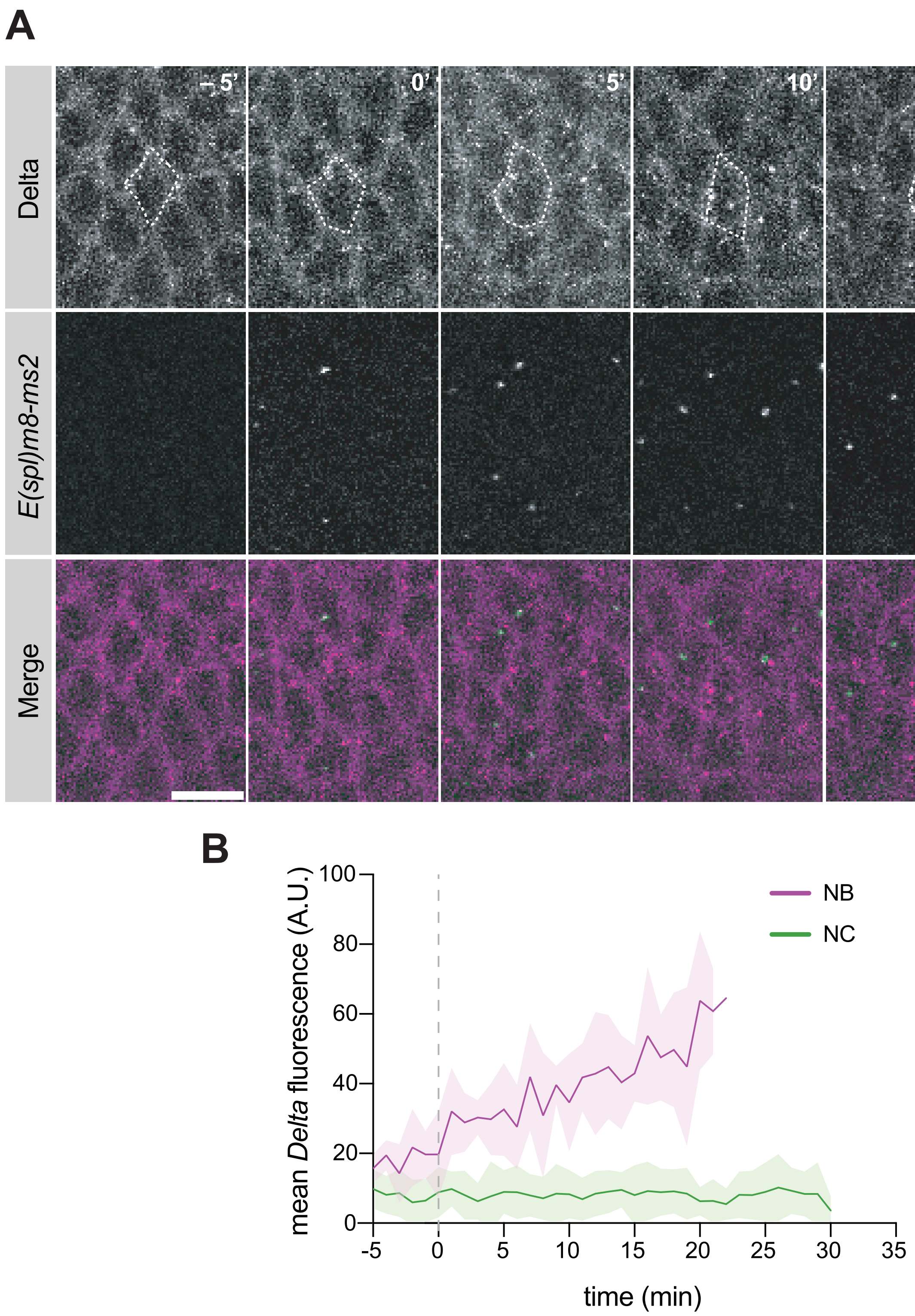
A
-5'
0'
5'
10'
20'
25'
Delta
E(spl)m8-ms2
Merge
B
mean Delta fluorescence (A.U.)
100
80
60
40
20
0
-5
0
5
10
15
20
25
30
35
time (min)
NB
NC

#### Supplementary Information: Additional details regarding computational model

To examine how cell geometry and mechanical properties influence signalling dynamics, we modified the lateral inhibition model described by[1], (which builds on the framework introduced by[2]. Unlike the original formulation, our implementation does not rely on a spatial activator gradient. Instead, we defined a perimeter-weighted cell–cell connectivity matrix, such that the interaction strength between neighbouring cells is proportional to the length of their shared boundary.

To account for possible mechanical influences on signalling behaviour, we introduced two additional parameters that vary inversely with cell perimeter. The first parameter, $\kappa_{tens}$, represents a tension-related scaling factor that regulates the amount of activated Delta available to engage in trans-activation of Notch receptors on adjacent cells. The second parameter, $\kappa_{cis}$, controls the strength of cis-inhibition between Notch and Delta within the same cell (Figure 6A).

These extensions allow cell size to directly influence signalling dynamics within the model framework. In the following section, we present the full derivation of the model based on the formulation of[1], incorporating the modifications described above.

##### Notation

$n_i$ ≡ Notch levels in cell i

$d_i$ ≡ Delta levels in cell i

$d_i^A$ ≡ Activated Delta in cell i

$S_i$ ≡ [NICD, Mam, Su(H)] complex levels in cell i

$E_i$ ≡ *E(spl)* levels in cell i

$A_i$ ≡ Activator levels in cell i

$\langle d_j \rangle_{\mathrm{i}}$ ≡ sum of Delta from cells j, on the boundaries with cell i

$\langle n_j \rangle_{\mathrm{i}}$ ≡ sum of Notch from cells j, on the boundaries with cell i

$[n_i d_j]$ ≡ complex of Notch from cell i and Delta from neighbour cell j

$p_i$ ≡ perimeter of cell i

The following Michaelis-Menten reaction scheme corresponds to the processes illustrated in Figure 6A:

##### Delta activation

$$d_i + A_i \xrightarrow{\alpha^+} d_i^A + A_i \tag{1}$$

$$d_i^A \xrightarrow{\alpha^-} d_i \tag{2}$$

with $\alpha^+$ and $\alpha^-$ the Delta activation and de-activation rates respectively.

**Trans-activation, signal and transcriptional feedback**

$$n_i + d_j \underset{\kappa^-}{\overset{\kappa^+}{\rightleftharpoons}} [n_i d_j] \tag{3}$$

$$n_i + d_j^A \underset{\kappa^-}{\overset{\kappa^+}{\rightleftharpoons}} [n_i d_j^A] \tag{4}$$

Assuming that only activated Delta can produce signal:

$$[n_i d_j^A] \xrightarrow{\kappa_{trans}} s_i \rightarrow E_i \dashv A_i \tag{5}$$

with $\kappa^+$ and $\kappa^-$ the association and dissociation between Notch, Delta and the Notch-Delta complex. $\kappa_{trans}$ is the trans-activation rate.

**Cis-inhibition**

Notch and Delta within the same cell can bind however the resulting complex does not lead to signal. Here we assume that the complex can either degrade or dissociate back to free Notch and Delta.

$$n_i + d_i \underset{\kappa_{cis}^{-non}}{\overset{\kappa_{cis}^{+non}}{\rightleftharpoons}} [n_i d_i] \xrightarrow{\kappa_{cis-deg}^{non}} \emptyset \tag{6}$$

$$n_i + d_i^A \underset{\kappa_{cis}^{-A}}{\overset{\kappa_{cis}^{+A}}{\rightleftharpoons}} [n_i d_i^A] \xrightarrow{\kappa_{cis-deg}^{A}} \emptyset \tag{7}$$

where $\kappa_{cis}^{+non}$ and $\kappa_{cis}^{+A}$ are association rates, $\kappa_{cis}^{-non}$ and $\kappa_{cis}^{-A}$ are dissociation rates while $\kappa_{cis-deg}^{non}$ and $\kappa_{cis-deg}^{A}$ are the degradation rates of cis-inhibited Delta-Notch complexes from non-activated and activated Delta respectively.

The reactions outlined above can be expressed as the following system of differential equations, using a Hill-function formulation:

**Differential equations**

$$\frac{dd_i}{dt} = \beta_d + \alpha^- d_i^A - \alpha^+ A_i d_i - \gamma_d d_i - \left(\kappa^+ \langle n_j \rangle_{\mathrm{i}} d_i - \kappa^- [\langle n_j \rangle_{\mathrm{i}} d_i]\right) - (\kappa_{cis}^{+non} n_i d_i - \kappa_{cis}^{-non} [n_i d_i]) \tag{8}$$

$$\frac{dd_i^A}{dt} = \alpha^+ A_i d_i - \alpha^- d_i^A - \gamma_d^A d_i^A - \left(\kappa^+ \langle n_j \rangle_{\mathrm{i}} d_i^A - \kappa^- \left[\langle n_j \rangle_{\mathrm{i}} d_i^A\right]\right) - (\kappa_{cis}^{+A} n_i d_i^A - \kappa_{cis}^{-A} [n_i d_i^A]) \quad (9)$$

$$\frac{dn_i}{dt} = \beta_n - \gamma_n n_i - \left(\left(\kappa^+ n_i \langle d_j \rangle_{\mathrm{i}} - \kappa^- \left[n_i \langle d_j \rangle_{\mathrm{i}}\right]\right) + \left(\kappa^+ n_i \langle d_j^A \rangle_{\mathrm{i}} - \kappa^- \left[n_i \langle d_j^A \rangle_{\mathrm{i}}\right]\right)\right) - ((\kappa_{cis}^{+non} n_i d_i - \kappa_{cis}^{-non} [n_i d_i]) + (\kappa_{cis}^{+A} n_i d_i^A - \kappa_{cis}^{-A} [n_i d_i^A])) \quad (10)$$

$$\frac{d\left[n_i \langle d_j \rangle_{\mathrm{i}}\right]}{dt} = \kappa^+ n_i \langle d_j \rangle_{\mathrm{i}} - \kappa^- \left[n_i \langle d_j \rangle_{\mathrm{i}}\right] \quad (11)$$

$$\frac{d\left[n_i \langle d_j^A \rangle_{\mathrm{i}}\right]}{dt} = \kappa^+ n_i \langle d_j^A \rangle_{\mathrm{i}} - \kappa^- \left[n_i \langle d_j^A \rangle_{\mathrm{i}}\right] - \kappa_{trans}^A \left[n_i \langle d_j^A \rangle_{\mathrm{i}}\right] \quad (12)$$

$$\frac{dS_i}{dt} = \kappa_{trans}^A \left[n_i \langle d_j^A \rangle_{\mathrm{i}}\right] - \gamma_s S_i \quad (13)$$

$$\frac{dE_i}{dt} = \frac{\beta_E \left(\frac{S_i}{T_s}\right)^{c_s}}{1 + \left(\frac{S_i}{T_s}\right)^{c_s}} - \gamma_E E_i \quad (14)$$

$$\frac{dA_i}{dt} = \frac{\beta_A}{1 + \left(\frac{E_i}{T_E}\right)^{c_E}} - \gamma_A A_i \quad (15)$$

$$\frac{d[n_i d_i]}{dt} = \kappa_{cis}^{+non} n_i d_i - \kappa_{cis}^{-non} [n_i d_i] - \kappa_{cis-deg}^{non} [n_i d_i] \quad (16)$$

$$\frac{d[n_i d_i^A]}{dt} = \kappa_{cis}^{+A} n_i d_i^A - \kappa_{cis}^{-A} [n_i d_i^A] - \kappa_{cis-deg}^{A} [n_i d_i^A] \quad (17)$$

$\beta_d, \beta_n, \beta_E, \beta_A$ are expression rates of Delta, Notch, *E(spl)* and activator respectively,

$\gamma_d, \gamma_n, \gamma_s, \gamma_E, \gamma_A$ are degradation rates of Delta, Notch, trans-activation signal, E(spl) and activator respectively,

$T_s$ and $T_E$ are the Hill half occupations levels associated with trans-activation signal promoting *E(spl)* expression and *E(spl)* inhibiting activator expression respectively, while $c_S$ and $c_E$ are the respective Hill coefficients.

Assuming that association rates of cis-inhibitory complexes are the same for all Delta variations:

Let $\kappa_{cis}^{-non} = \kappa_{cis}^{-A} = \kappa_{cis}^{-}$ , $\kappa_{cis}^{+non} = \kappa_{cis}^{+A} = \kappa_{cis}^{+}$ and $\kappa_{cis-deg}^{non} = \kappa_{cis-deg}^{A}$

Assuming steady-state for Delta-Notch & [NICD, Mam, Su(H)] complexes:

$$\frac{d\left[n_i\langle d_j\rangle_{\mathrm{i}}\right]}{dt}=0\ \Rightarrow\ \left[n_i\langle d_j\rangle_{\mathrm{i}}\right]=\frac{\kappa^+}{\kappa^-}\,n_i\langle d_j\rangle_{\mathrm{i}}\ ;\ \left[\langle n_j\rangle_{\mathrm{i}}d_i\right]=\frac{\kappa^+}{\kappa^-}\,\langle n_j\rangle_{\mathrm{i}}d_i \tag{18}$$

$$\frac{d\left[n_i\langle d_j^A\rangle_{\mathrm{i}}\right]}{dt}=0\ \Rightarrow\ \left[n_i\langle d_j^A\rangle_{\mathrm{i}}\right]=\frac{\kappa^+}{\kappa^-+\kappa_{trans}^A}\,n_i\langle d_j^A\rangle_{\mathrm{i}}; \tag{19}$$

$$\left[\langle n_j\rangle_{\mathrm{i}}d_i^A\right]=\frac{\kappa^+}{\kappa^-+\kappa_{trans}^A}\,\langle n_j\rangle_{\mathrm{i}}d_i^A \tag{20}$$

$$\frac{d[n_i d_i]}{dt}=0\ \Rightarrow\ [n_i d_i]=\frac{\kappa_{cis}^+}{\kappa_{cis}^-+\kappa_{cis\_deg}}\,n_i d_i \tag{21}$$

$$\frac{d[n_i d_i^A]}{dt}=0\ \Rightarrow\ [n_i d_i^A]=\frac{\kappa_{cis}^+}{\kappa_{cis}^-+\kappa_{cis\_deg}}\,n_i d_i^A \tag{22}$$

$$\frac{ds_i}{dt}=0\ \Rightarrow S_i=\frac{\kappa_{trans}^A}{\gamma_s}\left[n_i\langle d_j^A\rangle_{\mathrm{i}}\right] \tag{23}$$

$$\Rightarrow S_i=\frac{\kappa_{trans}^A}{\gamma_s}\frac{\kappa^+}{\kappa^-+\kappa_{trans}^A}\,n_i\langle d_j^A\rangle_{\mathrm{i}} \tag{24}$$

Let

$$K_t\equiv\frac{\kappa^+\kappa_{trans}^A}{\kappa^-+\kappa_{trans}^A} \tag{25}$$

be trans-activation rate and

$$K_c\equiv\frac{\kappa_{cis}^+\kappa_{cis\_deg}}{\kappa_{cis}^-+\kappa_{cis\_deg}} \tag{26}$$

be cis-inhibition rate.

New equations after the above assumptions and definitions:

$$\frac{dd_i}{dt}=\beta_d+\alpha^-d_i^A-\alpha^+A_id_i-(\gamma_d+K_cn_i)d_i \tag{27}$$

$$\frac{dd_i^A}{dt}=\alpha^+A_id_i-\left(\alpha^-+\gamma_d^A+K_t\langle n_j\rangle_{\mathrm{i}}+K_cn_i\right)d_i^A \tag{28}$$

$$\frac{dn_i}{dt}=\beta_n-\gamma_nn_i-K_tn_i\langle d_j^A\rangle_{\mathrm{i}}-K_c\left(d_i+d_i^A\right)n_i \tag{29}$$

$$\frac{dE_i}{dt} = \frac{\beta_E \left(\frac{K_t}{\gamma_s T_s}\right)^{c_s} \left(n_i \langle d_j^A \rangle_{\mathrm{i}}\right)^{c_s}}{1 + \left(\frac{K_t}{\gamma_s T_s}\right)^{c_s} \left(n_i \langle d_j^A \rangle_{\mathrm{i}}\right)^{c_s}} - \gamma_E E_i \quad (30)$$

$$\frac{dA_i}{dt} = \frac{\beta_N}{1 + \left(\frac{E_i}{T_E}\right)^{c_E}} - \gamma_A A_i \quad (31)$$

**Non-dimensionalization**

As in [1], we normalize all of the levels by the initial Notch level, $n_0$, and scale the time by the *E(spl)* degradation rate, $\gamma_E$, assuming degradation rates are equal among variables:

$$\gamma_d = \gamma_n = \gamma_s = \gamma_E = \gamma_A$$

Dimensionless equations after simplifications are:

$$\frac{dd_i}{dt} = \beta_d + \alpha^- d_i^A - \alpha^+ A_i d_i - (1 + K_c n_i) d_i \quad (32)$$

$$\frac{dd_i^A}{dt} = \alpha^+ A_i d_i - (\alpha^- + 1 + K_t \langle n_j \rangle_{\mathrm{i}} + K_c n_i) d_i^A \quad (33)$$

$$\frac{dn_i}{dt} = \beta_n - n_i - K_t n_i \langle d_j^A \rangle_{\mathrm{i}} - K_c \left(d_i + d_i^A\right) n_i \quad (34)$$

$$\frac{dE_i}{dt} = \frac{\beta_E \left(\frac{K_t}{T_s} n_i \langle d_j^A \rangle_{\mathrm{i}}\right)^{c_s}}{1 + \left(\frac{K_t}{T_s} n_i \langle d_j^A \rangle_{\mathrm{i}}\right)^{c_s}} - E_i \quad (35)$$

$$\frac{dA_i}{dt} = \frac{\beta_A}{1 + \left(\frac{E_i}{T_E}\right)^{c_E}} - A_i \quad (36)$$

**Introducing perimeters**

A perimeter is introduced for each cell and the connectivity matrix (which determines the function of Notch and Delta on boundaries between cells) is weighed using these perimeters.

The original connectivity matrix (in [1]):

$$C_{ij} = \begin{cases} \frac{1}{6}, & if\ i\ and\ j\ are\ neighbors \\ 0, & otherwise \end{cases} \quad (37)$$

is adapted using estimations of the contact lengths between cells using their perimeters:

$$L_{ij} = \frac{\frac{1}{2}(p_i + p_j)}{\langle \frac{1}{2}(p_i + p_j) \rangle_{\mathrm{C_{ij>0}}}} \tag{38}$$

$$C_{ij}^{w} = C_{ij} L_{ij} \tag{39}$$

Hypothesizing that cis-inhibition is inversely related to cell-size potentially reflecting membrane crowding effects that limit the availability of free Notch, we introduce perimeter dependent parameter $\kappa_{cis}$, as

$$\kappa_{\mathrm{cis,i}} = \left(\frac{\langle \mathrm{p} \rangle}{p_i}\right)^{\gamma_{cis}} \tag{40}$$

which scales the cis-inhibition factor as follows:

$$K_{c,i} = \kappa_{\mathrm{cis,i}} K_c \tag{41}$$

We also introduce a perimeter-based tension factor, $\kappa_{tens}$, based on the hypothesis that higher tension in smaller cells results in more activated Delta at their boundaries (for example via effects on endocytosis) available for trans-activation neighbouring Notch:

$$\kappa_{\mathrm{tens,i}} = \frac{\left(\frac{\langle \mathrm{p} \rangle}{\mathrm{p_i}}\right)^{\gamma_{tens}}}{\langle \left(\frac{\langle \mathrm{p} \rangle}{\mathrm{p_i}}\right)^{\gamma_{tens}} \rangle} \tag{42}$$

This factor re-scales the amount of activated Delta available for trans-activation as follows:

$$\langle d_j^A \rangle_{\mathrm{i}} = \sum_j C_{ij}^{w} \kappa_{\mathrm{tens,j}} d_j^A \tag{43}$$

For the non-dynamic, imposed perimeter model shown in Figure 6, we use the following perimeter rules:

(i) a smooth, time-dependent decrease in NB perimeter and

(ii) an increase in perimeter in cells whose *E(spl)* levels exceeded the *E(spl)* detection threshold as follows:

$$p_i(t) = \big(1 - s(t)\big)p_{0,i} + s(t)p_{target,i} \tag{44}$$

where,

$$s(t) = \begin{cases} 0 & t < t_0 \\ \dfrac{t - t_0}{\tau} & t_0 \leq t < t_0 + \tau \\ 1 & t \geq t_0 + \tau \end{cases} \tag{45}$$

and

$$p_{target,i} = \begin{cases} 0.05 & for\ NB \\ 1.4p_{0,i} & if\ E_i > \mathrm{L_D} \\ p_{0,i} & otherwise \end{cases} \tag{46}$$

where $p_i$is the perimeter of cell i (with $p_{0,i}$ the initial perimeter),

$\tau$ is the delamination duration and $t_0$ the timepoint when this rule is triggered

and $\mathrm{L_D}$ is *E(spl)* detection limit.

To test whether our model would work with fully dynamic perimeter behaviour rather than imposed changes (Figure S4), we introduce a perimeter non-dimensionalised ordinary differential equation. As before, time is scaled by the *E(spl)* degradation rate, $\gamma_E$. Perimeter evolution is represented as a function of intracellular *E(spl)* and activated Delta levels using a Hill function formulation:

$$\frac{dp_i}{dt} = -\kappa_p \frac{d_{A,i}^h}{\theta_\delta^h + d_{A,i}^h} + \frac{E_i^m}{\theta_E^m + E_i^m} \tag{47}$$

where $\kappa_p$ is the ratio of Activated Delta and *E(spl)* Hill function prefactors, $\theta_\delta$ and $\theta_E$ are the Hill activation thresholds associated with Delta activity inhibiting perimeter and *E(spl)* promoting perimeter respectively, while $h$ and $m$ and the respective Hill coefficients.

#### Computer simulation

The dimensionless equations are solved numerically, using a standard Matlab ODE solver, and the solution is simulated on a 8x8 2D hexagonal lattice (Figure 6D) using a periodic boundaries condition.

For the model with imposed perimeter changes (Figure 6A) the following success criteria are used:

(i) *E(spl)* expression occurred only in NCs, with at least one NC exceeding the detection threshold and

(ii) the ratio between maximal *E(spl)* levels in NCs and the detection threshold exceeded 10, consistent with calibration of our MS2 system[3].

For model with dynamic perimeter (Figure S4) two additional conditions are added:

(i) NB perimeter had to decrease below 0.3 by the end of the simulation, and

(ii) the mean perimeter of non-NB, non-NC cells had to remain above 0.6.

Following our non-dimensionalization procedure, as described above, the time in the simulation is in units of the typical half-life of key proteins in the system (estimated to be ~2.5mins)[3].

The following parameters are used, with parameters from the original model retaining the values reported in that study (see [1] for justification):

**Table S1: Initial values and parameters for model**

| Parameter | Description | Value |
|---|---|---|
| $\beta_d$ | Delta production rate | 1.62 |
| $\beta_n$ | Notch production rate | 1.52 |
| $\beta_E$ | *E(spl)* production rate | 1.62 |
| $\beta_A$ | Activator production rate | 1.62 |
| $\alpha^+$ | Delta activation rate | 0.6 |
| $\alpha^-$ | Delta de-activation rate | 0.4 |
| $K_t$ | Trans-activation rate | 0.8 |
| $K_c$ | Cis-inhibition rate | 0.4 |
| $T_E$ | Threshold level for *E(spl)* to inhibit activator expression | 0.0051 |
| $T_s$ | Threshold level for trans-activation signal to promote *E(spl)* expression | 0.8 |
| $c_s$ | Hill coefficient for promoting *E(spl)* expression by trans-activation signal | 3 |
| $\gamma_{tens}$ | Exponent scaling "tension" factor $\kappa_{tens}$ | 3 for Figures 6D,F and Figure S4 |
| $\gamma_{cis}$ | Exponent scaling cis-inhibition factor $\kappa_{cis}$ | 1 for Figures 6D,F<br>0 for Figure S4 |
| $t_0$ | Simulation timepoint that perimeter rule (44) is activated | 0 |
| $\tau$ | Delamination duration associated with perimeter rule (44) | 10 for Figures 6B-E,<br>[5, 10, 15] for Figure 6F |
| $L_D$ | *E(spl)* detection threshold | [0.010 0.013 0.016 0.019] for Figures 6B,C |

| | | |
|---|---|---|
| | | 0.015 for Figures 6D,F<br>[0.015 0.02 0.025] for Figures S4A,B<br>0.025 for Figure S4C,D |
| $\kappa_p$ | Ratio of Activated Delta and *E(spl)* Hill function prefactors in dynamic perimeter ODE (47) | [0.15, 0.20, 0.25] for Figures S4A, B<br>0.2 for Figures S4C,D |
| $\theta_\delta$ | Threshold level for Delta activity to inhibit perimeter | 0.4 |
| $\theta_E$ | Threshold level for *E(spl)* expression to promote perimeter | 0.4 |
| $h$ | Hill coefficient for inhibiting perimeter by Delta activity | 2 |
| $m$ | Hill coefficient for promoting perimeter by *E(spl)* expression | 1 |
| $d_0$ | Initial levels of Delta | 1 ± 5% noise |
| $d_0^A$ | Initial levels of activated Delta | 0 |
| $n_0$ | Initial levels of activated Notch | 1 ± 5% noise |
| $E_0$ | Initial levels of *E(spl)* | 0 |
| $A_0$ | Initial levels of activator | 0.6 ± 5% noise |
| $p_0$ | Initial perimeter values | 0.85 for presumptive NB, 1 ± 5% noise for all other cells for Figures 6B-E<br>0.75 for presumptive NB, 1 ± 5% noise for all other cells for Figures S4C,D |
